# Transcriptome-wide profiling of acute stress induced changes in ribosome occupancy level using external standards

**DOI:** 10.1101/2022.05.30.493937

**Authors:** Annie W. Shieh, Sandeep K. Bansal, Zhen Zuo, Sidney H. Wang

**Affiliations:** Center for Human Genetics, The Brown foundation Institute of Molecular Medicine, The University of Texas Health Science Center at Houston, Houston, TX; Baylor College of Medicine, Houston, TX

**Author notes:** These authors contributed equally to this work (Annie W. Shieh, Sandeep K. Bansal). **coauthor email:**, AWS, SKB, ZZ.

## Abstract

Acute cellular stress is known to induce a global reduction in protein translation through suppression of cap dependent translation. However, selective translation in response to acute stress has been shown to play important roles in regulating the stress response. An accurate transcriptome-wide profile of acute cellular stress-induced translational changes has been challenging to obtain. Commonly used data normalization methods, such as quantile normalization, operate based on the assumption that any systematic shifts are artifacts introduced from experimental procedures. Consequently, if applied to profiling acute cellular stress-induced protein translation changes, these methods are expected to produce biased estimates. To address this issue, here we designed, generated, and evaluated a panel of 16 oligomers to serve as external standards for ribosome profiling studies. Using Sodium Arsenite treatment-induced oxidative stress in lymphoblastoid cell lines as a model system, we applied spike-in oligomers as external standards based on quantifications of monosomal RNA extracted from each sample. We found our spike-in oligomers to display a linear correlation between the observed and the expected, with small but significant ratio compression at the lower concentration range, and span the expected quantitative range in the observed data, which covers 97 % of the quantitated endogenous genes. We found popular global scaling normalization approaches to introduce both high levels of false positives and false negatives in differential expression analysis. Using the expected fold changes constructed from spike-in external controls, we found in our dataset that TMM normalization produced 87.5% false positives when a P value cutoff of 0.1 is used (i.e. 10% expected false positive rate)% and on average produced a systematic shift of fold change by 3.25 fold. These results highlight the consequences of applying global scaling approaches to conditions that clearly violate their key assumptions. As an alternative, we found using spike-in quantifications as control genes in RUVg normalization recapitulated the expected stress induced global reduction of translation and resulted in little, if any, systematic shifts in spike-in constructed true positives. Finally, using spike-in constructed true positives and true negatives, we explored alternative normalization approaches for acute cellular stress response ribo-seq studies. We found that a simple approach that quantile normalized data from control and treated samples separately, which we termed respective quantile normalization, produced expected results in spike-in quantification, and resulted in little, if any, systematic bias on fold change in endogenous genes. Additionally, we found that under certain parameters, using endogenous control genes for RUVg normalization best recapitulate the expected. Our results clearly demonstrated the utility of our spike-in oligomers, both for constructing expected results as controls and for data normalization. Our exploration of different normalization approaches highlights the issues in applying global scaling normalization when key assumptions are clearly not met. We show that a respective quantile normalization approach or normalization with endogenous control genes are viable options worth considering as more generalizable approaches for stress response ribo-seq studies. This conclusion is likely applicable to other types of studies that involve global shifts in expression profiles between comparison groups of interests.

## Introduction

Advances in high throughput gene expression profiling technologies, such as microarrays and next generation sequencing technologies, have revealed many technical challenges in making seemingly straightforward comparisons. Oftentimes, despite great care in implementing experimental procedures, systematic shifts in quantification range are observed between samples, and sometimes even between samples that are otherwise considered identical. In response to these technical challenges, many procedures have been developed to identify and to remove these unwanted variations ^1–9^. Approaches such as standardizing read count by library size and global scaling to homogenize quantitative range across samples are among the most commonly used strategies for increasing statistical power to detect differential expression and to improve accuracy in effect size estimates ^2,5^. These powerful approaches, however, make certain assumptions that need to be met in order to deliver the intended results^10^. For example, a global scaling normalization approach, such as TMM, assumes that any systematic shift in quantitative range is an experimental artifact, and scales the data accordingly to mitigate the systematic bias. Such assumption is often valid, as a clear global shift would indicate an unlikely scenario that most, if not all, genes are changing expression level in the same direction relative to the control samples/conditions. In some scenarios, however, a systematic shift in gene expression is a part of the biological process of interest. For example, a key step of the Unfolded Protein Response (also known as Integrated Stress Response) is the phosphorylation of a translation initiation factor eIF2-alpha, which in turn shuts down cap-dependent translation in order to reduce protein synthesis load in the ER^11–15^. In such a scenario, a global reduction in protein translation is expected. Applying global scaling normalization on a dataset profiling stress induced changes in protein translation level will therefore erroneously normalize away many of the stress induced changes by severely distorting the effect size of most, if not all, quantitated genes.

In a scenario where a global reduction in gene expression is expected, a normalization approach alternative to global scaling is needed to accurately quantitate gene expression. External standards designed for quality control to facilitate cross platform comparison, such as ERCC RNA spike-in control mixes^16,17^, could be used in such experiments to provide reference points for normalization. A straightforward regression based standard curve approach could connect estimates of gene expression level to RNA concentration, however, this approach is not suitable for normalization efforts to increase power in differential expression tests. Using libraries constructed with the same ERCC spike-in pool, Jiang et al. found significant deviations in the observed fold change from the expected, despite high Pearson correlation for spike-in quantification between libraries^17^. The observed deviations exceeded two fold for spike-in controls in the lower abundance range, likely arising simply from sampling variation^17^. Risso et al. contemplated the use of spike-in controls for data normalization and demonstrated potential issues with spike-in based global scaling approaches and a spike-in based local regression approach ^1,2,18^. Instead, Risso et al. proposed a factor analysis approach for control gene based normalization (RUVg) by treating quantifications of spike-in oligomers as control genes. Although Risso et al. found spike-in oligomers clearly violated a key assumption of their method, i.e. spike-in oligomers should recapitulate the reaction to unwanted effects by endogenous genes, they found the method to generate expected results and outperformed its contemporaries^1^.

While ERCC spike-in oligos are available for RNA-Seq studies, they are not easily adaptable to ribosome profiling experiments, an approach enabling transcriptome-wide profiling of protein translation^19^. Consequently, transcriptome-wide profiling of translation level changes in response to external stimuli, especially for those that have the potential to induce global shift in quantification, has been challenging^20^. Earlier stress response ribo-seq studies therefore opted for either not normalizing the data or normalizing the data with available methods that were not necessarily suited for the purpose ^21,22^. On the other hand, Andreev et al. used a single oligomer spike-in as external control for data standardization in their study on the impact of Sodium Arsenite treatment (oxidative stress) on protein translation in HEK293T cells^23^. Their analysis, however, was limited by a rather small sample size and by their design of a single spike-in oligomer control, which does not enable more rigorous assessments on normalization impact. Work from Iwasaki et al. and subsequently Liu et al used mitochondrial ribosome footprints as controls for normalization and achieved apparent success ^24,25^. However, as was articulated by the authors, the underlying assumptions supporting the use of mitochondrial footprints as controls for normalization may or may not be met in different experimental conditions. Such an approach is therefore unlikely to be generalizable. Even in studies where the results appear to meet expectations, the caveat of observing false results introduced by potential systematic shifts in mitochondrial ribosome footprints could undermine the conclusion.

To meet this apparent need, a few recent studies have explored the utility of different spike-in formulations as external standards for data normalization^26,27^. Here we report our efforts on this front. We developed and further characterized a panel of 16 spike-in control oligomers designed for ribosome profiling experiments. To characterize these spike-in oligomers and demonstrate their utility in ribo-seq data normalization, we conducted a study to profile transcriptome-wide changes in ribosome occupancy level induced by Sodium Arsenite treatment. Our spike-in formulation and study design provides the expected fold changes between samples, which enables rigorous evaluation of normalization impact on effect size and false positive rate.

## Results

### A set of short spike-in control oligos designed for human ribo-seq experiments

Our spike-in oligos design follows Lutzmayor et al. oligo design for small RNA-Seq^28^. The key features are 1. A core sequence at position 5-24 that mimics the base composition of the first 20 nts of human miRNA. 2. Four-nucleotide-random-sequences flanking each end. 3. A free energy profile mimicking endogenous miRNA. 4. End modifications to mimic ribosome footprints (Fig. S1a). Using this design, we generated 993 core sequences each in combination with 65,536 different permutations of flanking sequences. We selected 16 oligo sequences, one from each category of core sequence, based on their free energy profile similarity to the endogenous miRNA (Fig. S1b); 16 different spike-in oligomers were ordered from the manufacturer in two batches of 8 (Table S1). These synthetic oligos were then mixed to create spike-in pools following the modified Latin square design by Pine et al.^29^ to create pools of spike-in oligos that cover a relative quantitative range of 1∼17,920 (Fig. 1a). We apply these spike-in pools for ribo-seq experiments by adding the oligo mix to the RNA sample prepared from the monosome isolation step using size exclusion column, which we termed monosomal RNA. Because the majority of monosomal RNA are not ribosome footprints, through trial and error, we empirically determined the ratio between spike-in oligo pool and monosomal RNA that will result in a desirable amount of spike-in oligomers sequenced in the final ribo-seq library. We found that, in our system, a 1:20,000 weight ratio between the spike-in pool and the monosomal RNA results in an average of 2.30 % final spike-in count/ total footprint count in the sequencing data.

**Figure 1.**
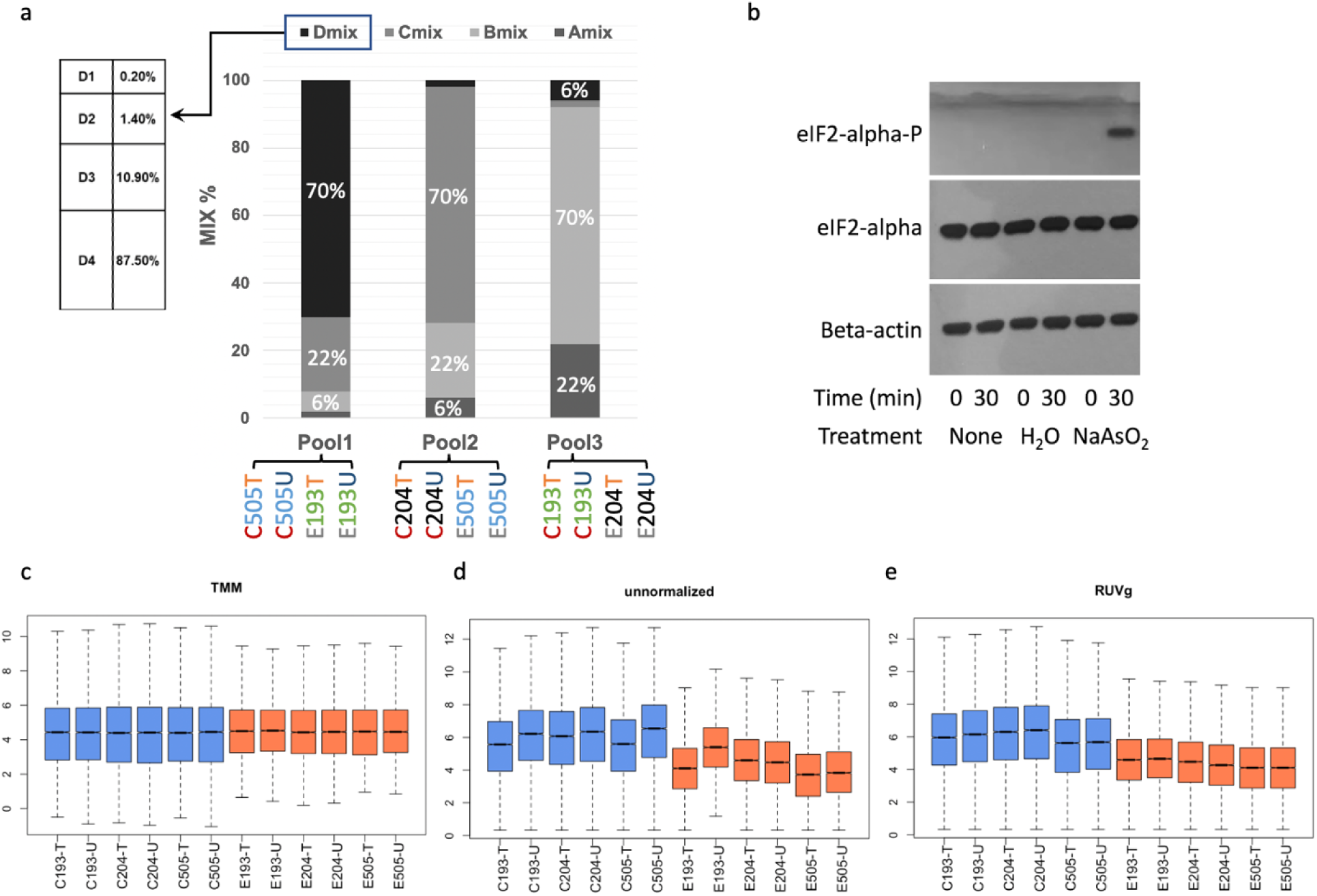
The design and application of spike-in oligomers for stress response ribo-seq studies. (a) Spike-in oligomer pooling design: A stacked barplot showing proportion of spike-in mixes used in each spike-in pool. Each mix is composed of 4 spike-in oligomers (e.g. D1, D2, D3, D4) mixed in the same 8 fold increment from oligomer 1 to oligomer 4 (proportion in percentages presented in the table to the left for mix D). Below the stacked barplot, samples receiving each spike-in pool are labeled with color code distinguishing different effects of interests to highlight the study design. Sodium Arsenite treatment (Control vs. Experiment). Donor (18505 vs. 19204 vs. 19193). Library preparation batch (T vs. U). (b) Western blots for GM18505 indicate that our treatment conditions induced integrated stress response. Primary antibodies used were labeled to the left of the blots. Treatment types and durations were labeled below the blots. Treatment type “None” indicated baseline conditions, while treatment type “H_2_O” indicated the control condition used in the ribo-seq study. Note the band of phosphorylated eIF2α, a stress marker, only visible in Sodium Arsenite treated samples in contrast to the loading controls. (c, d, e) Boxplots summarizing impact of different normalization strategies on overall distribution of ribosome occupancy level across all genes analyzed. Boxes are color coded to distinguish control samples (blue) from Sodium Arsenite treated samples (orange). The maximum and minimum values for the boxplot (i.e. the whiskers) are defined by the genes with quantification levels closest to (but without exceeding) 1.5 times of the interquartile range extending from the box.

### A stress response ribo-seq study for evaluating spike-in utility

We aim to evaluate the utility of spike-in oligos in a scenario where a global shift in translation level is expected. As such, we collected ribo-seq data with spike-in from three unrelated HapMap LCL cell lines (GM18505, GM19193, GM19204) with and without 30 minutes of Sodium Arsenite induced Oxidative stress. A 30 minute treatment with Sodium Arsenite at a final concentration of 134 μM is sufficient to induce phosphorylation of eIF2α in LCL (Fig. 1b). Phosphorylation of eIF2α is a hallmark of integrated stress response and is expected to result in a global reduction of cap-dependent protein translation ^11,12^. From each sample we created 2 technical replicates of sequencing libraries, one of which was treated with an additional CRISPR depletion step aimed to further reduce the level of rRNA contamination in ribo-seq libraries. We generated a total of 873 million sequencing reads, which, after filtering out the rRNA, tRNA, and snoRNA reads, resulted in an average of ∼2 million ribosome footprint reads uniquely mapped to the GRCh38 human genome and ∼32K reads mapped to the spike-in oligo sequences (Table S2). We found no significant differences in the proportion of rRNA reads between libraries with and libraries without the CRISPR depletion step (P=0.82, Wilcoxon rank sum). To focus our analysis on sufficiently quantitated genes, we required, for each GENCODE annotated gene, at least one sequencing read mapped to the gene in each sample of our dataset. With this criteria our sequencing coverage enabled analysis of 12,357 genes. We found that the panel of 16 spike-in oligomers span similar quantitative ranges across samples, which on average covers 97% of the quantitated genes (Fig. S2a). Without further normalization, we readily observed the expected global reduction in ribosome footprint counts across quantitated genes in the Sodium Arsenite treated samples (Fig. S2a, Fig. 1d).

### Quality assessment for spike-in oligos

We first evaluate the relationship between spike-in sequencing counts and their corresponding nominal concentrations. Overall we observe the expected strong positive correlations across samples with median Spearman’s rho at 0.87, ranging between 0.81 to 0.92 (Fig. S2b). We found no significant effect from treatment conditions (P = 0.86, ANOVA) nor from technical replication batches (P = 0.19, ANOVA) on the aforementioned relationship. On the other hand, we found the oligos from two separate orders to show clear ordering batch effects (Fig. S2c; P= 1.43e-4, ANOVA). Although it is unclear what manufacturing process drives this batch effect, it is clearly important to consider the batch effect from manufacturing, especially if the oligos were intended for use in a standard curve. We next evaluate the property of individual spike-in oligos in order to identify outlier oligos that are potentially not suitable for use as external controls. We first evaluate the monotonicity of oligo quantification. While experimental conditions could introduce variations in quantification between samples, oligo proportions within a sample are not expected to be affected by these external factors. The oligo quantifications are therefore expected to follow a monotonic function determined by the pooling fraction (i.e. the order of quantification should correspond to the order of fraction in the pools). Using Spearman’s Rho, we found all spike-in oligos used in this study are monotonic. To observe the impact of concentration range and outlier data points, for each spike-in oligo, we evaluate the Pearson correlation between the expected and the observed quantification level. Overall We found strong correlations between the expected and the observed (median 0.94, interquartile range 0.92∼0.97). On the other hand, for spike-in oligos used in the lower concentration range, we observed a decrease in correlation between the observed and the expected (Fig. 2a). Of note, we found oligo A2_752 to be an outlier of the observed trend (Fig. 2a). A Pearson correlation between the observed and the expected for A2_752 is 0.54, which fell far below the 95% confidence interval of a loess fit adjusting for the expected concentration range from each spike-in. Next, we evaluate the expected fold change for each spike-in oligomer between samples based on our modified Latin square pooling design (Fig. 1a). Overall, we found a small but significant ratio compression effect in the observed fold change relative to the expected (Fig. 2b). When using the expected fold change of spike-in oligomers in a linear model to predict the observed fold change, we found a regression slope of 0.87 +/-0.023, which is significantly different from the expected regression slope of 1 (P = 2.28e-6). This ratio compression effect appears to be stronger when oligomers in the lower concentration range are used (Fig. 2c). By stratifying data into three groups of the concentration range covered in our study, we found a progressive increase in ratio compression in the lower concentration group (Fig. S3), which points to the possibility of an additive process such as consistent pipetting bias during serial dilution as the cause of the observed ratio compression effect. This observed significant ratio compression effect, albeit small, points to additional technical challenges in using spike-in oligos for a standard curve.

**Figure 2.**
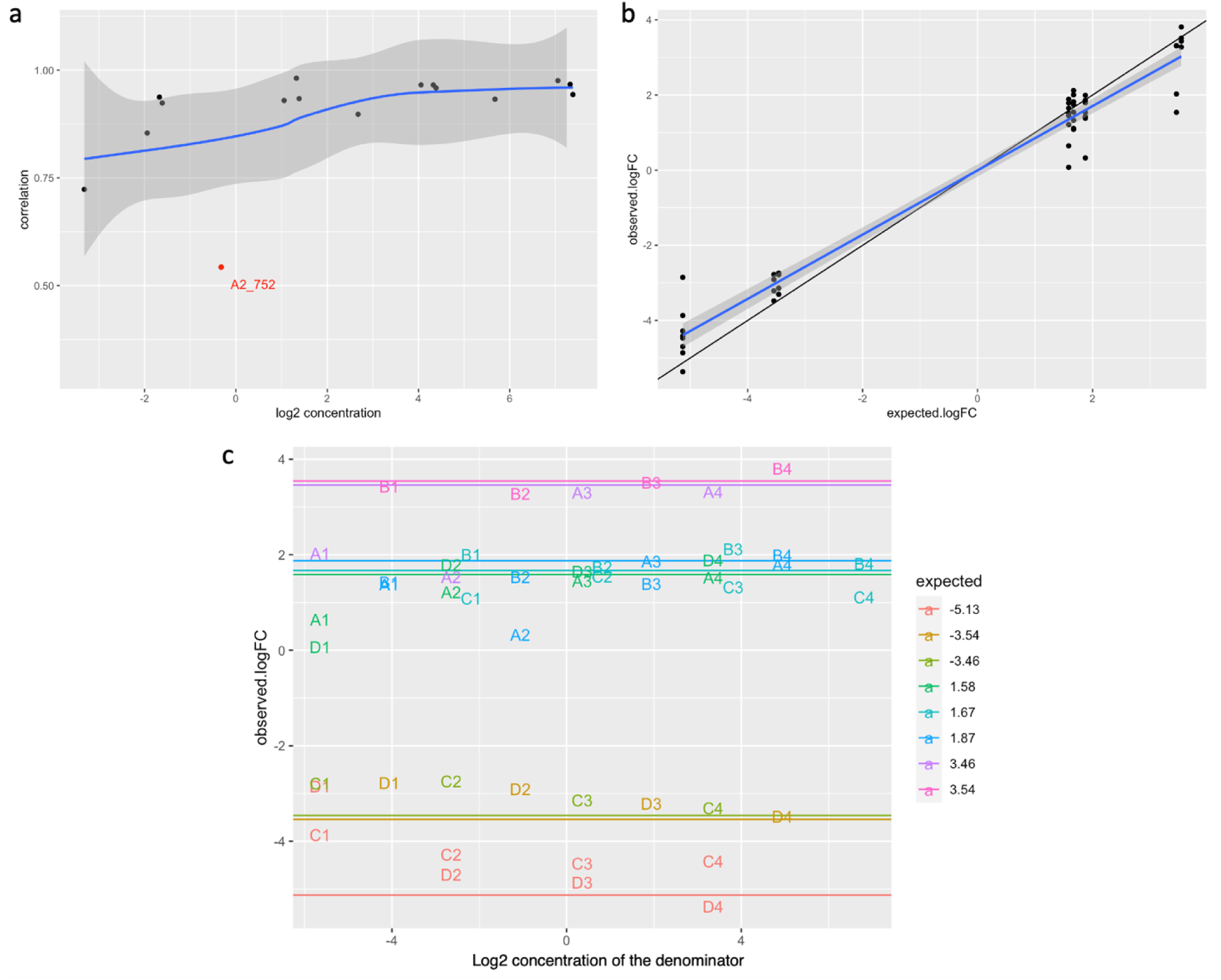
Quantitative properties of Spike-in oligomers. (a) Correlations between the observed quantification and expected for each spike-in oligomer is plotted against the across-pool-average-expected concentration of the spike-in oligomer. Note the downward trend towards the lower concentration range. Blue loess trend line and its corresponding 95% confidence interval (shaded area) indicated oligomer A2 (highlighted in red) as an outlier. (b) Observed log2 fold change for spike-in constructed true positives plotted against the expected. Each data point represents the fold difference of the same oligomer between 2 different spike-in pools. The blue trendline and shaded area represents the linear regression coefficients and corresponding 95% confidence intervals calculated using the expected log fold change as predictor. (c) Observed log2 fold change for spike-in constructed true positives plotted against the expected concentrations of the quantifications used as the denominator for the observed log2 fold change calculation. The horizontal lines mark the expected log2 fold change. Each data point is represented by corresponding oligomer identification abbreviations and color coded in the same way as the expected fold change lines. Note the ratio compression at the lower concentration range.

### Spike-in based data normalization preserved the expected global shift in quantification

It has previously been shown that a standard curve normalization approach using control oligos can be technically challenging^17^. Consistently, evaluation of our spike-in oligos has shown both manufacturing batch effect and ratio compression effect that render a standard curve approach using our spike-in oligomers impractical. Alternatively, Risso et al. has developed a factor analysis approach - RUV, which could use spike-in oligos as “control genes” for normalizing RNA-seq data^1^. Here we apply RUVg to our ribo-seq stress response dataset and compare the results with those that were generated from applying a popular global scaling normalization approach, TMM^2^, on the same dataset. The first step of applying RUVg for data normalization is to determine the number of unwanted variables (k) to remove. To this end, we evaluate the impact of removing unwanted variables on the correlation between the observed spike-in quantification and the expected relative concentration in conjunction with the available degrees of freedom afforded by the sample size of our dataset. By incrementally removing unwanted variables, we found that increasing correlation between the observed and the expected spike-in quantification started to plateau at k =3. This choice of k is robust against the exact composition of spike-in oligomer quantification data used for normalization (Fig S4). We therefore decided on removing the top 3 unwanted variables, which increased the correlation between the observed spike-in quantification and the expected from R^2^ = 0.73 to 0.83. Using spike-in based RUVg normalization, we found the resulting normalized dataset to preserve the expected global reduction in ribosome occupancy level (Fig. 1e). In contrast, when applying TMM normalization to the same dataset, the expected global shift is lost (Fig. 1c).

The expected concentration of spike-in oligomers can be used to construct true positives with expected fold change and true negatives in differential expression tests. Using the set of true negatives, we compare the number of false positive discoveries between unnormalized data, TMM normalized data, and RUVg normalized data. At a P value cutoff of 0.1, we found ∼20% false positives from tests using either unnormalized data or RUVg normalized data. Conversely, at the same cutoff using TMM normalization, we found 42 false positives out of a total of 48 tests (87.5%). The extremely high proportion of false positives identified using TMM normalized data is in clear contrast to the proportion of false positives identified either using unnormalized data or RUVg normalized data, both consistently mere ∼10% higher than the expected proportion defined by the p value cutoffs (Fig. S5). On the other hand, when comparing the expected and observed fold change for true positives constructed based on differences in spike-in proportions between spike-in pools, we found TMM normalization to clearly distort the observed fold change (Fig. 3a). Fold change estimates calculated based on TMM normalized spike-in oligomer quantifications deviate from the expected (regardless of the direction of effect) by 68% (+/-6.3%) of the expected fold change. In contrast, 31% (+/-4.2%) and 34% (+/-3.8%) of absolute deviation from the expected fold change were respectively observed for unnormalized and RUVg normalized sipke-in quantifications. The more than doubled absolute deviation observed in TMM normalized data (i.e. relative to the unnormalized) appears to be mainly driven by the TMM normalization procedure shifting the quantification in treated samples upwards. By modeling the observed log2 fold change in TMM normalized spike-in quantification using the expected log2 fold change as the predictor, we found TMM normalization to maintain a regression coefficient of 0.824 +/-0.038 and a r-squared of 0.910, both values comparable to the unnormalized data (regression coefficient: 0.832 +/-0.033; r-squared: 0.930). On the other hand, when comparing between spike-in oligomer quantifications in the direction of treatment relative to the control samples, we found the log2 fold change of TMM normalized spike-in quantification significantly deviates from the expected by 1.612 +/-0.143 (a systematic upward shift, P < 2e-16); in contrast, −0.082 +/-0.129 log2 deviations from the expected fold change were observed for unnormalized data (Fig. S6). The distortion in fold change estimates observed in TMM normalized data explains the high false positive rates observed in the aforementioned true negative control comparisons and indicates the possibility of introducing false negatives by the global scaling normalization approach shifting true differences towards 0. On the other hand, RUVg normalization appears to have nudged the observed fold change towards the expected (Fig. 3a), which resulted in an increase in the regression coefficient, when using the expected log2 fold change as the predictor, from 0.832 (+/-0.033) to 0.882 (+/-0.041) and a change in the estimated intercept from −0.082 (+/-0.105) to 0.011 (+/-0.128), both values inching closer to the expected.

**Figure 3.**
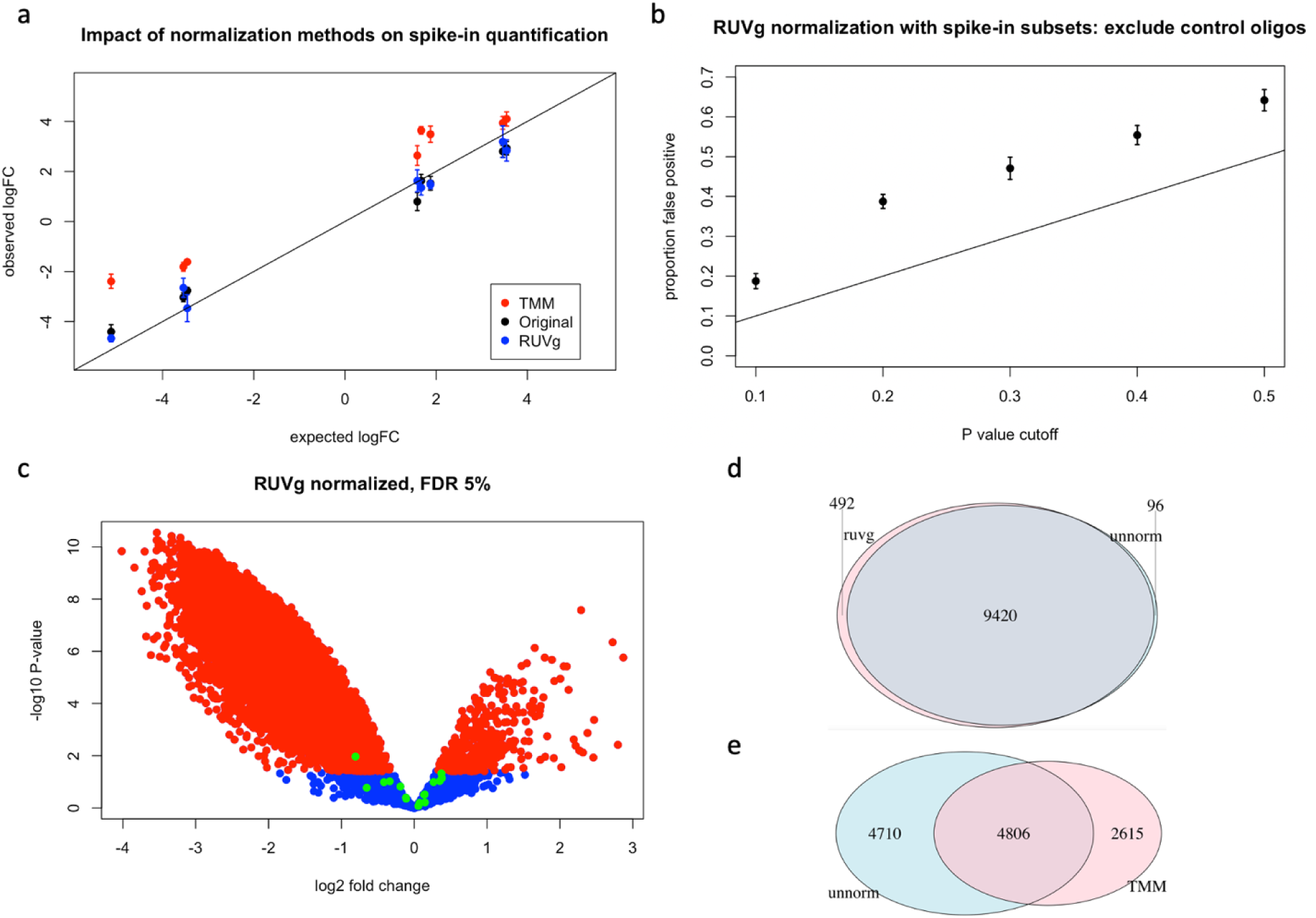
Impact of TMM normalization and spike-in based RUVg normalization on expression quantification. (a) Observed log2 fold change for spike-in constructed true positives plotted against the expected; comparing between results from different normalization approaches. Data points represent mean plus/minus standard errors calculated from each group of true positives that share the same expected log2 fold change. Black data points (i.e. “Original”) are results from log2 transformed counts without further normalization. Black line indicates the perfect correlation (i.e. an intercept of 0 and a slope of 1). (b) Proportion of significant differences (i.e. false positives) found in spike-in constructed null comparisons across a range of P value cutoffs. Error bars represent the standard error across 10 iterations of random subsampling of spike-in quantifications as control genes for RUVg normalization. The black line marks the expected number of false positives. (c) A volcano plot showing the relationship between fold change (treatment versus control) and p value from differential expression tests for RUVg normalized data. Data points are color-coded in green for spike-in oligomers, in red for endogenous genes that are significantly differentially expressed at 5% FDR, and in blue for endogenous genes that are not differentially expressed. (d, e) Venn diagrams summarizing the number of shared and distinct differentially expressed genes found with and without normalization. Unnormalized (unnorm) in blue. Normalized (either TMM or RUVg) in pink. Numbers labeling each area indicate the number of genes belonging to each group, with the size of the area drawn in proportion to the size of the group.

To evaluate the robustness of spike-in-based RUVg normalization, we took a subsampling approach. For each iteration we randomly subsampled half of our spike-in quantification data as control genes. We compared RUVg normalization results across ten iterations of subsampling. For the true negatives, we found the distribution of fold change to consistently center near the expected 0 across iterations (−0.016 +/-0.027, Fig. S7a). In addition, similar to the results from normalizing using the full set of spike-in data, we found the proportion of false positives to consistently fall ∼10% above the expected across a wide range of significance cutoffs (Fig. S7b). Qualitatively similar, albeit noisier, results were observed when analyzing only spike-in quantification data that were not included as control genes for normalization (fold change: −0.021 +/-0.045, proportion false positives: Fig. 3b). For the spike-in comparisons designed to have expected fold changes (i.e. true positives) we found the observed fold changes to have a tight range of values across subsampling iterations for both the absolute deviations from the expected (33.8% +/-0.3% of the expected) and the parameters of a linear model using expected log fold change as the predictor (regression coefficients: 0.888 +/-0.002, intercept:-0.016 +/-0.027), each set of values covers the respective estimates from using the full set of spike-in quantification data for normalization (Fig. S8).

After evaluating the impact of normalization procedures on spike-in quantification, we next compare RUVg normalized gene quantification results to results from TMM normalization. At 5% FDR we identified 9,912 stress induced translation level changes from 12,357 genes using spike-in based RUVg normalized data. Consistent with the observed global reduction in translation activities, 96% of stress induced changes were down regulations (Fig. 3c). When compared with test results from unnormalized data, RUVg normalization replicated 99% of the discoveries (i.e. 9,420 out of 9,516) and identified an additional 492 genes (∼5%) (Fig. 3d). In contrast, using TMM normalization, at 5% FDR we identified 7,421 stress induced translation level changes from 12,357 genes and of these differentially translated genes only 48% of stress induced changes were down regulations (Fig. S9b). When compared with test results from unnormalized data, TMM normalization only replicated 51% of the discoveries (i.e. 4,806 out of 9,516) yet identified an additional 2,615 genes (i.e. 35% of TMM discoveries) (Fig. 3e). The high proportion of TMM only discoveries is consistent with the high false positive rates observed in the spike-in negative control analysis. In addition, the low replication rate on the differential expression detected in unnormalized data (i.e. 51% compared to the 99% from RUVg normalization) indicates a potential high false negative rate resulting from TMM normalization. To further evaluate the possibility of normalization induced false positive discoveries, we contemplate the possibility of recovering differentially expressed genes found only in normalized data by relaxing significance cutoff in differential expression tests applied to unnormalized data. A relaxed significance cutoff is expected to recover some of the true differences that were missed simply because of power issues. For differentially expressed genes found in TMM normalized data but not in unnormalized data (at 5% FDR), we found limited recovery of differentially expressed genes by relaxing significance cutoffs (Fig. S10a). Using 20% FDR, we recovered only 25.6% of the normalization specific findings. Moreover, amongst the recovered genes 69.4% had an opposite direction of effect, indicating a high rate of false recoveries. In contrast, for RUVg normalization, we found a much higher recovery rate (Fig. S10b). Using 20% FDR, we recovered 86% of the normalization specific findings and all of the recovered effects were found in the same direction as the unnormalized data. To evaluate the possibility of TMM normalization generating high levels of false negative results from our dataset, we focus our analysis on the differentially translated genes identified from unnormalized data that were not replicated in TMM normalized data. We found that TMM normalization shifted the observed log2 fold change in treated samples relative to control sample upwards by 1.733 +/-0.001, which resulted in the TMM normalized log2 fold change to center near 0 (Fig. S10c). This upward shift is comparable in magnitude to the upward shift observed in spike-in oligomers (1.612 +/-0.143) (Fig. S6), indicating the same underlying cause pushing the observed fold change upwards in spike-in oligomers could be producing false negatives results in genes by shifting the observed effect size towards zero. In other words, in our dataset, TMM normalization appears to produce false negatives mainly by shifting effect size as opposed to introducing noise. In fact, when comparing log2 fold change between TMM normalized and unnormalized data across all quantitated genes, it becomes apparent that TMM normalization shifts the entire distribution of log2 fold change (Fig. S10d), which, judging from its impact on spike-in quantification, led to inaccurate fold change estimates for all genes, and as a result produced both false positives and false negatives. Taken together, these results highlight the issues in using a global scaling approach, such as TMM, to normalize ribo-seq datasets in scenarios of a global shift in protein translation, such as acute cellular stress response. Such scenarios also highlight the utility of external standards for data normalization.

### Evaluating alternative normalization approaches using spike-in control oligomers

Mitochondrial ribosome footprints have previously been used for normalizing stress response ribo-seq data by standardizing gene counts with per sample total mitochondria footprint counts^24,25^. This approach postulates that being in a separate cellular compartment, translation events in the mitochondria are not (immediately) subjected to stress-induced eIF2α phosphorylation and the consequent global reduction of protein translation. In such a scenario, ribosome footprints from mitochondrial transcripts could be used as controls to standardize cytosolic ribosome footprints. Using spike-in oligomers, we sought to evaluate this postulate in our experimental condition. Before normalization, we found no clear systematic shifts between treatment and control samples in the log transformed mitochondrial footprint counts, albeit a high level of inter-sample variation in count distribution is observed (Fig. S11). To normalize cytosolic ribosome footprint counts with mitochondrial footprints, we use a “mitoTMM” approach, in which the total mitochondrial footprint counts were used to calculate an effective library size via the TMM method, and this effective library size was then used for cytosolic ribosome footprint count normalization. Overall, we found the global shift in cytosolic translation in response to oxidative stress is preserved in mitoTMM normalize data (Fig. 4a). We next evaluate the impact of mitoTMM on spike-in quantification. When evaluating true negative comparisons (i.e. comparing between samples derived from different donors that received the same pool of spike-in oligos) we found a high level of false positives (Fig. S12a). At a P value cutoff of 0.1, we found 24 false positives out of 48 true negative comparisons (50%). In contrast, when evaluating the expected fold change (i.e. comparing between spike-in pools in samples derived from the same donor) we found a strong positive correlation between the observed fold change and the expected with little effect size distortion (regression coefficient: 0.85, R^2^: 0.91; Fig. S12b). By visualizing the effect of mitoTMM normalization on the observed fold change in scatterplots (Fig. S12c,d), we found that, for true negatives, systematic trimmed-mean shifting of count distribution produced false positive results by shifting the observed fold change away from zero with a relatively modest effect size (∼2 fold). On the other hand, trimmed mean shifting had a more muted effect on true positive comparisons that resulted in a limited number of false negatives and an overall distribution similar to the unnormalized (Fig. S12d). A combination of the larger expected effect sizes for true positives and a more balanced systematic trimmed-mean shifting explains the attenuated impact. Because fold change shifting by trimmed mean centering is based on the effective library size calculated using mitochondrial footprint counts from each sample, the samples involved in each expected versus observed comparison determines the amount of distribution shift. In our design, the by-pool-comparison group for the true negatives is confounded by the donor identity. In other words, the high false positive rate observed could be resulted from TMM shifting distribution to counteract the donor effects. Alternatively, given the small sample size for these control comparisons (n=4), it is possible that the observed difference in false positive rates between mitoTMM normalized data and the unnormalized data simply reflects fluctuations in sampling variation. When making true negative comparisons using a larger sample size (n= 12) and a different comparison group design that mimics differential expression tests for genes (i.e. by reordering and comparing across different oligomers of the same expected concentration between the control and the treated samples), we found no false positives out of 16 true negative comparisons at either the 5% FDR cutoff used for differential expression tests across the transcriptome (Fig. 4b) or a P value cutoff of 0.1 (the smallest P value is 0.17). The same could not be said for data transformed by a standard TMM normalization approach using total ribo-seq counts for calculating effective library size, in which 7 out of 16 true negative comparisons were found significantly differentially expressed at 5% FDR (Fig. S9b), which corresponds to 10 false positives out of 16 null comparisons (i.e. 62.5%) at a P value cutoff of 0.1. Using mitoTMM normalization, we identified 9,676 stress induced translation level differences from 12,357 genes at 5% FDR. Similar to results from using spike-in RUVg normalization, 96% of the observed differential expression events were down regulation in the treated samples. When compared with test results from unnormalized data, mitoTMM normalization replicated 98% of the discoveries (i.e. 9,324 out of 9,516) and identified an additional 352 genes (∼4%). On the other hand, when comparing the observed log2 fold change between mitoTMM normalized data and unnormalized data (Fig. S13), we found a slight systematic upward shift in mitoTMM normalized data (0.1 +/-0.0004, p < 2e-16), indicating potential false positives and false negatives for genes with small effect size.

**Figure 4.**
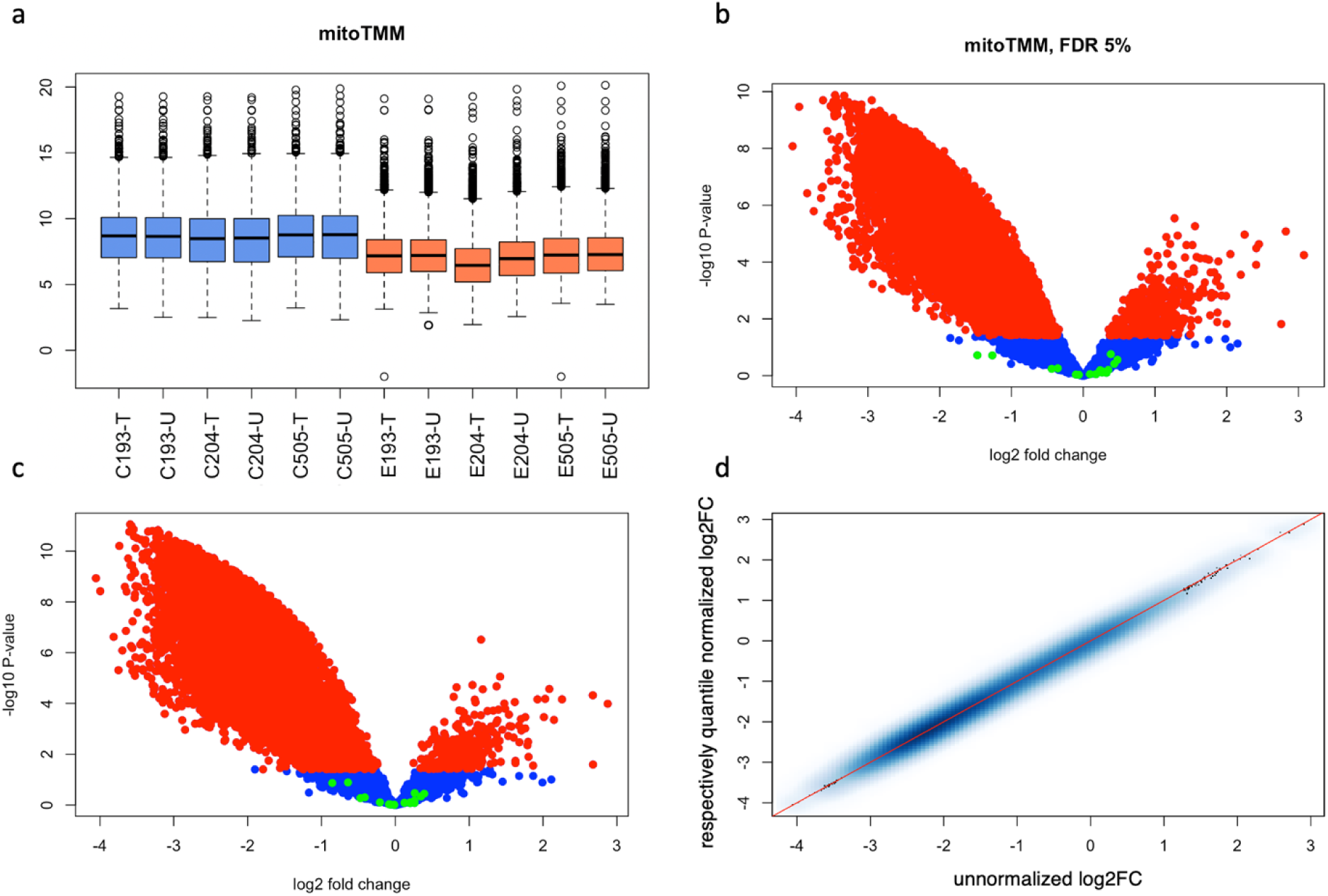
Using spike-in quantifications as controls to evaluate alternative normalization approaches. (a) Boxplots summarizing overall distributions of expression quantifications for endogenous genes across samples after mitoTMM normalization. Boxes are color-coded to distinguish control samples (blue) from Sodium Arsenite treated samples (orange). The maximum and minimum values for the boxplot (i.e. the whiskers) are defined by the genes with quantification levels closest to (but without exceeding) 1.5 times of the interquartile range extending from the box. (b, c) Volcano plots showing the relationship between fold change (treatment versus control) and p value from differential expression tests for mitoTMM normalized data (b) and for respectively quantile normalized data (c). Data points are color-coded in green for spike-in oligomers, in red for endogenous genes that are significantly differentially expressed at 5% FDR, and in blue for endogenous genes that are not differentially expressed. (d) Log2 fold changes between treatment and control samples for respectively quantile normalized data plotted against the unnormalized in a smoothed color density representation of a scatter plot. All endogenous genes deemed sufficiently quantitated are included in the plot with color density proportional to the number of genes presented in each unit region (bin) of the plot. Red line marks the perfect correlation.

The apparent effectiveness of the mitoTMM approach prompted us to evaluate a more generalizable global scaling approach for normalizing stress response ribo-seq data. To this end, we considered a respective global scaling normalization approach, in which control samples and treated samples were separately normalized to their respective average distributions. A respective quantile normalization approach is counter intuitive, because it confounds the “normalization batch” with the effect of interest, and therefore risks amplifying systematic biases. On the other hand, it remains unknown whether such bias exists in our dataset. Using spike-in control oligos, we sought to evaluate the impact of respective quantile normalization on differential expression tests. As expected, respective quantile normalization preserves the global shift in the level of protein translation in treated samples (Fig. S14a); on the other hand, no systematic differences were observed in spike-in oligomer quantification between control and treated samples (Fig. S14b). When evaluating spike-in constructed true negatives and true positives, we found the results similar to mitoTMM normalization. For true negatives, we found 25 significant differences out of 48 null comparisons (52.1%) at a P value cutoff of 0.1. Note that although the numbers of false positive discoveries are similar between mitoTMM and respective quantile normalization, different sets of comparisons are found to be false positives (Fig. S14c). When using an alternative comparison group design that evaluates true negatives across all samples, we found no significant differences at either the 5% FDR cutoff used for differential expression tests across the transcriptome (Fig. 4c) or a P value cutoff of 0.1 (the smallest P value is 0.13). For true positives, we again found a strong positive correlation between the observed fold change and the expected with little effect size distortion (regression coefficient: 0.82, R^2^: 0.93; Fig. S14d). Using respective quantile normalization, we identified 9,797 stress induced translation level changes from 12,357 genes at 5% FDR. Consistent with a global reduction in protein translation in response to oxidative stress, 97% of the observed differential expression events were downregulated in the treated samples. When compared with test results from unnormalized data, respective quantile normalization replicated 99% of the discoveries (i.e. 9,374 out of 9,516) and identified an additional 423 genes (∼4%). Of note, when comparing the observed stress treatment effect between respectively quantile normalized data and unnormalized data, we found little to no systematic shifts (Fig. 4d), indicating limited, if any, systematic biases were introduced by separately normalizing the control samples from the treated samples.

The spike-in based RUVg normalization approach suffers from two major caveats: 1. Only a very limited number of control genes were available for estimating unwanted effects. 2. Spike-in oligomers are not known to behave similarly to endogenous genes in response to unwanted effects, which is an explicit assumption of the RUVg method^1^. An alternative use of the RUVg method by employing endogenous control genes could side-step both caveats. However, without prior knowledge, it is challenging to identify control genes. Using spike-in oligomers, we aim to evaluate the impact of endogenous control gene selection on RUVg normalization results. We first define overlapping sets of control genes by using a series of P value cutoffs for differential expression tests between treatment and control samples in unnormalized data; we anticipate the list of genes that failed to reach significance cutoffs to be enriched of genes that have no real differences between conditions. Regardless of the choice of P value cutoffs for defining the control geneset, the correlation between the observed and the expected quantification for spike-in oligomers plateaued at around R^2^ = 0.83 after 5 unwanted variables were removed (i.e. k = 5). A lack of variation in linear model fit between the observed and the expected across different control gene sets indicates the possibility that RUVg normalization removed relevant biological effects that distinguish between different control gene sets. On the other hand, we found the incremental sets of control genes appear to have variable impacts on normalization results when either 3 or 4 unwanted variables were removed (i.e. k = 3 or k = 4) (Figure 5a). Among the control genesets, we found the set defined by P value > 0.7 to result in the highest R2 of 0.82 between the observed and the expected in data normalized using K=4 RUVg approach. To further evaluate the differential impact between removing 4 and 5 unwanted variables (i.e. k=4 vs. k=5), we compared normalization impact on differential expression test results using the same control gene set define by genes with P value > 0.7 in differential expression tests performed on unnormalized data comparing between treated and control samples. For data normalized with k = 5 RUVg approach, we found the normalization to increase both the number of differentially expressed genes identified and the number of false positive discoveries from spike-in constructed null comparisons. At 5% FDR, differential expression tests on k = 5 RUVg normalized data identified 10440 differentially translated genes (an increase of 924 from 9516) and identified 3 false positives from 16 null comparisons. Using a P value cutoff of 0.1 instead of the 5% FDR used, we found 7 false positives from 16 null comparisons (44%), which far exceeded the expected 10% false positive rates for null comparisons. In contrast, for data normalized with k = 4 RUVg approach, at 5% FDR we found RUVg normalization to recover 99.57 % of differentially expressed genes found in unnormalized data, and further increased the number of differentially expressed genes identified from 9516 to 10234, without introducing false positive results in spike-in null comparisons (Figure 5b). In this case, using a P value cutoff of 0.1 instead of 5% FDR, we found 3 false positives from 16 null comparisons (19%), which is about twice the expected 10%; note that at a P value cutoff of 0.05, only 1 false positive was identified from 16 null comparisons, which correspond to a false positive rate of 0.0625, which is on par with the expected value of 0.05. Consistent with the results from null comparisons made across all samples, for data normalized using k = 4 RUVg approach, results from within pool comparisons for spike-in constructed true negatives consistently fall near the line of the expected false positive proportions (Fig. 5c), while results from between pool comparisons for spike-in constructed true positives show an increase in the regression coefficient from 0.832 (+/-0.033) of the unnormalized to 0.899 (+/-0.044) and a change in the estimated intercept from −0.082 (+/-0.105) to −0.055 (+/-0.138), both values moving closer to the expected.

**Figure 5.**
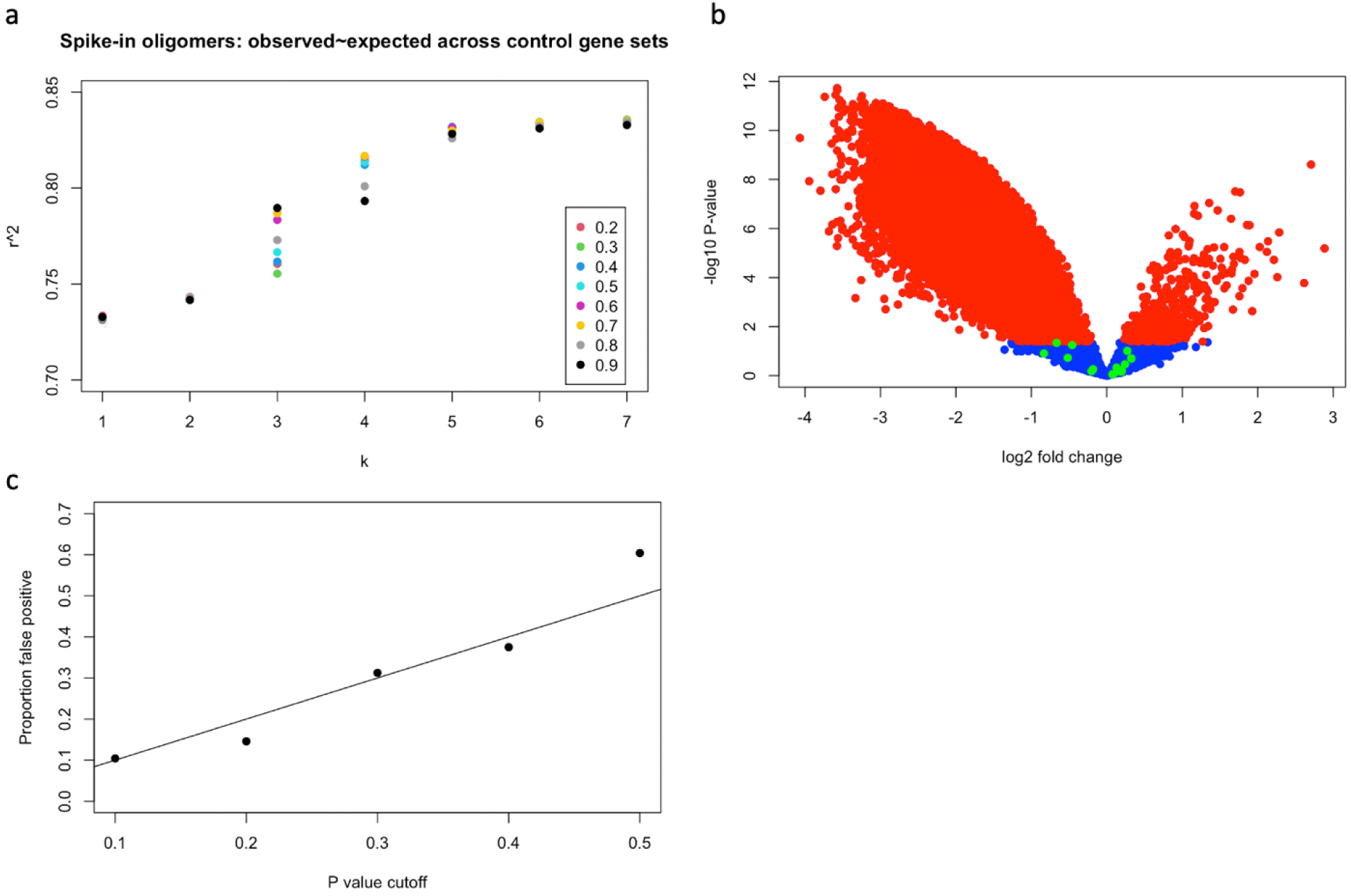
Using spike-in quantifications to evaluate results from endogenous-control-gene-based RUVg normalization. (a) Impacts of the number of unwanted variables removed and the control gene set used are visualized in a scatterplot of the correlation between the observed and the expected for spike-in quantifications plotted against the number of unwanted variables removed. Data points are color coded by the P value cutoff used to define control gene sets. A cutoff of 0.2 indicates that all genes have a P value greater than 0.2 from a test between control and treated samples in the unnormalized data are included as control genes. (b) A volcano plot visualizing the differential expression test results from RUVg normalized data using a control gene set defined by P > 0.7 and removing 4 unwanted variables (k = 4). Green data points are spike-in quantifications, blue and red data points indicate endogenous genes that pass the significance cutoff of 5% FDR. (c) Proportion of false positives identified from spike-in constructed null comparisons in endogenous-control-gene-based RUVg (P > 0.7, k = 4) normalized data. The black line indicates the proportion of false positives expected by chance.

## Discussion

We developed a panel of 16 spike-in oligomers and a corresponding pooling scheme for applications in ribosome profiling studies to identify acute cellular stress induced changes in translation levels across the transcriptome. To evaluate the utility of this set of spike-in oligos, we performed ribosome profiling experiments to identify Sodium Arsenite induced changes in translation level from 3 LCLs in 2 conditions each with 2 library preparation replicates. Evaluation of the quantitative properties of spike-in oligomers overall found strong positive correlation between the observed and the expected. At the same time, we identified one oligomer, A2, as potentially problematic and revealed two major unwanted effects, a manufacturing batch effect and a ratio compression effect. Because of the limited number of oligomers designed and the fact that our design requires a consistent length and has a rather homogenous base composition across oligomers, we were unable to identify length bias or sequence features associated with unwanted effects. On the other hand, our spike-in oligomers were designed to span a spectrum of minimum free energy (i.e. a numeric proxy for oligo folding structure) that resembles endogenous miRNAs, which allows us to evaluate the correlations between minimum free energy and the quantitative properties of the spike-in oligomers. We found no significant association between oligomer minimum free energy and the correlation between the observed and the expected oligomer quantification (P=0.60, Fig. S15). Of note, we found the outlier A2 oligomer to have the highest minimum free energy of 0 among the 16 spike-in oligomers. We designed the spike-in oligomers based on endogenous miRNA minimum free energy profile, because we frequently observed “ribosome protected fragments” pileup at miRNA loci. While it remains unclear what proportion of these miRNA fragments are true ribosome footprints and what proportion are resulted from miRNA RISC complex co-purified with monosomes during size exclusion column purification, these miRNA fragments clearly made their way into the ribo-seq data. We therefore reason that artificial oligomers mimicking miRNA folding and base composition will make good candidates as spike-in controls for ribo-seq studies. Although our results doesn’t allow us to evaluate whether a certain level of minimum free energy is required to ensure incorperations of the spike-in oligomers, a lack of correlation observed between spike-in performance and minimum free energy is reassuring, in that quantification level is not biased by the minimum free energy of RNA fragments. Based on the observed pattern of manufacturing batch effect, we postulate that between batch variations in either the quantification process from the manufacturer or variations in preparing the stock solution when we first received the oligomers (e.g. different sets of pipetman used or pipetman calibration shifted overtime) was the likely culprit. The progressive nature of ratio compression led us to postulate that a consistent pipetting error could have resulted in the observed compression, which is more pronounced at the lower concentration range.

Despite the unwanted effects observed, our spike-in oligomers, when used as control oligos, clarified the consequences of using a global scaling normalization approach to transform ribo-seq data collected from an acute cellular stress response study. When evaluating the expected true negatives constructed based on our spike-in pooling design, we found a high proportion of false discoveries in TMM normalized data, while in contrast, in data without normalization we found the proportion of false discoveries fell consistently below the FDR cutoffs used. When evaluating the expected true positives, we found TMM normalization to shift fold changes based on differences in sequencing coverage between libraries. In our Sodium Arsenite treatment study, such shifts ended up creating both false positives and false negatives. The troubling results observed from TMM normalization does not indicate problematic behaviors of TMM normalization method per se, instead, it reflects the consequences of violating an important underlying assumption for all global scaling normalization approaches, i.e. any systematic shifts are assumed to be experimental artifacts.

We explored the possibility of using spike-in oligomers for data normalization. Following work from Risso et al. we used a factor analysis approach, RUVg, to perform spike-in based data normalization. RUVg has previously been used for spike-in based normalization^1^. Despite violating a key assumption of the RUVg normalization method, i.e. control genes respond to unwanted effects similarly to the genes of interest, RUVg normalization using ERCC spike-in as control genes appeared to perform well^1^. Applying RUVg normalization to our stress response ribo-seq dataset using our 16 spike-in oligomers as control genes appears to have preserved the expected global reduction in protein translation in response to stress treatment and slightly increased power. Although the power increase is rather limited (because of the large effect size from stress treatment resulted in the majority of differentially expressed genes readily detectable without normalization), RUVg normalization could have increased accuracy in fold change estimates, as was indicated by the increase in regression coefficient (i.e approaching the expected value of 1), which was observed consistently across subsampling iterations. The oligomer manufacturing batch effect and ratio compression effect, although randomized across the effect of interest, are clearly not properties shared by endogenous genes. How these properties impact the parameter estimate for unwanted effects and therefore the normalization results remain to be determined. As a proxy for unknown unwanted effects, we visualized the impact of library preparation batch effect comparing between spike-in oligomers and endogenous genes (Fig. S16). Overall, we found a rather similar trend between spike-in oligomers and endogenous genes, with the exception of a bump in the middle of the distribution (Fig. S16), which could potentially be attributed to sampling variation; however, by the same token, the limited number of spike-in oligomers reduced confidence in our interpretation.

The caveats in using spike-in oligomers as control genes for RUVg normalization prompted us to explore alternative approaches for normalizing stress response ribo-seq datasets. Following previous publications ^24,25^, we evaluated the results from using mitochondria footprint for normalization. We found Sodium Arsenite treatment did not result in a systematic shift in mitochondrial ribosome footprints, and consequently, normalization using the mitoTMM approach appears to work reasonably well, with a relatively small systematic bias (Fig. S13). On the other hand, it is unclear how generalizable this conclusion is. Our experiment only treated cells with Sodium Arsenite for 30 mins, in an attempt to limit downstream impact on transcription^23^. It is unclear if a longer exposure or if using a different stimuli all together would result in a global shift in mitochondrial ribosome footprints. As a potentially more generalizable alternative, we evaluated the results from a respective quantile normalization approach, in which data from control and treated samples were first quantile normalized separately to their respective average distributions before combined together for differential expression tests. A respective normalization approach runs the risk of confounding the normalization batch and the effect of interest and potentially introducing systematic bias. However, our results indicated the approach to perform well in all metrics evaluated, including checking for systematic effect size shifts (Fig. 4d), except for the high false positive instances observed in the spike-in null comparisons (i.e. within-pool-between individual comparisons for the same spike-in oligomer). Similar high level of false positives in spike-in null comparisons was also observed in mitoTMM normalized data. In both cases, the false positives appear to be resulted from variation in sequencing coverages (a key piece of information heavily used by global scaling normalization methods), which is correlated with cell line identification in our dataset. In contrast, when using an alternative construction of spike-in null comparisons, which includes all samples, we found no false positives in respectively quantile normalized data. The clear decrease in false positive rate likely resulted from the increased noise from randomizing individual effects in the alternative construction of spike-in null comparisons. All things considered, it would appear that when the dataset is properly randomized and spiked with sufficient control oligos for quality check, a respective quantile normalization approach should be considered as a viable option for stress response ribo-seq studies.

Finally we evaluated an alternative RUVg normalization approach that uses endogenous control genes. By assuming that the unwanted effects on average introduce noises in the data as opposed to introducing biases (i.e. in regard to the stress treatment effect), we use a series of P value cutoffs to select overlapping sets of non-differentially expressed genes between control and treated samples in unnormalized data as control genes. Our analysis indicated that variations in normalization using different control gene sets are most apparent when either 3 or 4 unwanted variables were removed. Since we anticipated some control gene sets to work better than others (e.g. higher enrichment of true non-DE genes, more data points for estimating unwanted effects, etc.), we expected these variations to be informative. Indeed, by selecting the control gene sets that resulted in the highest correlation between the observed and the expected spike-in quantification when 4 unwanted variables were removed, we found the resulting dataset to produce differential expression test results that are well-calibrated. Wherein, the observed false positive proportions track closely with the expected, which is not seen in results from other normalization approaches. In contrast, when normalizing with removing 5 unwanted variables, the factors distinguishing between different control gene sets were removed by the normalization procedure and a high false positive rate in spike-in constructed null comparisons was observed. An increase in false positive proportion by removing an additional variable appears to be counterintuitive, because linear regression removes signals and is more commonly associated with false negatives. Given the construction of spike-in null comparisons, a likely explanation is that removing the additional variable led to a lower within-group-variation (i.e. lower variation between library replicates or between individual samples), which could result in smaller P values without increasing effect size for minor unadjusted unwanted effects. Whether such decrease in sampling variation also introduces false positives in endogenous genes requires further investigation. Given the small sample size, decreasing sampling variation without corresponding adjustment of minor unwanted effects could inadvertently introduce false positive results. In other words, our results indicated that there could be a healthy ratio between sampling variation and minor unwanted effects to avoid false positives. Further studies with more biological replicates, more technical replicates, and specifically designed spike-in pooling are needed to clarify this idea.

While our study aims specifically to address the unmet need of external standards for transcriptome-wide profiling of translational regulation in stress response, the exceedingly high proportion of false positives observed as a result of applying TMM normalized to our oxidative stress response ribo-seq dataset and the almost ubiquitous use of global scaling normalization approaches in the field of genomics prompts the unsettling question of how common such error might have occurred without the researchers acknowledging it. It is easy to envision scenarios with subtler global shifts in gene expression for studies comparing between developmental stages or testing for drug treatment effects. Without appropriate use of external standards, these subtle shifts could escape researchers and the consequent application of global scaling normalization will lead to distortion of effect size, which, as we have shown, results in both false positives and false negatives. While an appropriate application of external standards could safeguard against such pitfalls, it is not always obvious how much spike-in oligomers should be applied to each sample. With stress response ribo-seq studies, ample prior literature provided strong support for the use of total monosomal RNA (i.e. a proxy of ribosome number) as the reference point. Such information, however, is data type and biological context specific. Without a clear reference point for determining the appropriate amount of spike-in oligomers to use for each sample, large sample size will be needed to detect a subtle global shift. Considering the economy of research, a collective consciousness in including external spike-in oligomers in smaller scale studies and future retrospective meta analysis could provide a reasonable path forward.

## Materials and Methods

### Cell Culture and oxidative stress

Three lymphoblastoid cell lines (LCLs) (GM18505, GM19193, and GM19204), each derived from a separate Yoruba people from Nigeria, were purchased from Coriell Institute for Medical Research (NIGMS Human Genetic Cell Repository). The cells were maintained at 37°C with 5% CO_2_ in RPMI media supplemented with 15% FBS, 2 mM L-glutamate, 100 IU/ml penicillin, and 100 μg/ml streptomycin, in accordance with instructions provided by Coriell. Of note, cell cultures were vigilantly maintained at a cell density between 600,000 to 700,000 cells/ml to avoid inadvertent induction of stress response. To induce oxidative stress, cells were treated with 134 μM (i.e. the final concentration in cell culture) Sodium Arsenite (NaAsO_2_; Sigma-Aldrich, Cat # S7400), a heavy metal oxidative stressor, for 30 minutes in otherwise the same cell culture conditions. For the control group, nuclease free water was used in place of the Sodium Arsenite solution. After treatment, cells were pelleted by centrifugation at 100g for 10 min and washed twice with cold PBS (4°C). Cell pellets were flash frozen in liquid nitrogen and stored at −80°C.

### Western blot

Proteins were prepared from flash frozen cell pellets using M-PER Mammalian Protein Extraction Reagent (Thermo Scientific, Cat # 78503) following vendor’s instructions. Protein concentration was estimated using bicinchoninic acid (BCA) assay (Pierce™ BCA Protein Assay Kit, Cat# 23227) and 5 μg of proteins from each sample were used for western blot. All antibodies (e.g. anti eIF2α, anti Phospho-eIF2α) used were purchased from Cell Signaling Technologies (Table S2). XCell II Blot module was used for wet electrophoretic transfer of proteins from SDS-PAGE gel to nitrocellulose membrane (Biorad, Cat # 1620145, 0.45 μm) in Tris-Glycine Electroblotting buffer (National Diagnostic, Cat # EC-880). ProtoBlock Solution (National Diagnostics, Catalog number: CL-252) was used for blocking the nitrocellulose membrane following manufacturer’s instructions. After wet transfer, membranes were incubated with ProtoBlock solution for 1 hour at room temperature before overnight incubation with primary antibodies at 4°C (in blocking solution). After primary antibody incubation, membranes were washed in TBST buffer for three times (5 minutes each) before incubation with HRP conjugated secondary antibody for 2 hours at room temperature (in blocking solution). After secondary antibody incubation, membranes were washed three times in TBST buffer for 5 minutes each. ProtoGlow ECL (National Diagnostics, Cat # CL-300) and HyBlot CL autoradiography films (Thomas Scientific, Cat # NC1601219) were used for signal detection.

### Spike-in oligomers

Spike-in oligomer sequences were designed following Lutzmayer et al.^28^. While we followed the design principle and the programming scripts developed by Lutzmayer et al. a few key aspects were modified to suit our purpose of using these oligomers as external standards in ribosome profiling experiments for human samples. Instead of mimicking the free energy profile and base composition of *Arabidopsis* miRNA, our design mimics human miRNAs. In addition, we extended the oligomer length to 28 nts with flanking random tetramers to mimic ribosome footprint length. We generated/selected 1000 permutations of 20 nts RNA sequences that have a base composition resembling human miRNA (calculated based on a high confidence set of 896 miRNA sequences downloaded from miRBase^30^). Seven of these permutation sequences mapped to the human genome and were therefore removed. For the remaining 993 permutations of RNA sequences, we added random tetramers to each end of the sequence, which resulted in 993 sets of 65,536 sequences. Using RNAfold^31^, we determined the minimum free energies of all sequences. Based on the minimum free energy profile, we selected a total of 16 sequences (Table S1), each from a different set, to produce spike-in oligomers for the current study. These 16 spike-in oligomers were purchased in two separate batches (Table S1) from Sigma Aldrich and resuspended in 10 mM Tris (pH 8) to a stock concentration of 100 μM based on the quantifications provided by the vendor. To create the final spike-in pools used in experiments, four spike-in mixes (A, B, C, D) each composed of 4 different oligomers (1,2,3,4) in eight-fold concentration increments were created separately through 2-fold serial dilutions. The four mixes (A, B, C, D) were then combined in a defined ratio of (2%,6%,22%,70%) in 3 permutations to create 3 different spike-in pools each have individual oligomers in different concentrations but together covering the same concentration range (Fig. 1a). The working solution for spike-in pools was prepared in a weight concentration of 50 pg/μl, which we found convenient for an application of 50 pg spike-in pool for each 1μg of monosomal RNA before gel isolating the ribosome footprints.

### Ribosome profiling

Ribosome footprint profiling experiments were performed following the ligation free protocol described in Hornstein et al.^32^ with a few specific modifications made for the current study. Key steps include, RNase I digestion to generate ribosome protected fragments (100 U per 200 μl of cell lysate), size exclusion spin column (Sephacryl S400, GE: 27-5140-01) isolation of monosomes, spike-in oligomer control addition, gel isolation of ribosome footprints, sequencing library construction using Clontech SMARTER smRNA SEQ KIT (Fisher Scientific: NC1098027), rRNA depletion using subtraction oligo^33^, and finally PCR amplified Indexed libraries were pooled to sequence on an Illumina HiSeq 4000. For incorporating spike-in oligomers, for each sample, 50 pg of spike-in pool was used for each 1μg of monosomal RNA.

### CRISPR/Cas9 targeted depletion of rRNA

CRISPR/Cas9 mediated rRNA removal was performed following Han et al.^34^ and Mito et al.^35^. Ten target-specific oligomers for rRNA (see sequences in Han et al.^34^) were ordered from Sigma Aldrich and separately PCR amplified to prepare each oligomer as the template for in vitro transcription (T7-Scribe™ Standard RNA IVT, CELLSCRIPT, Cat# C-AS3107) to produce guide RNAs. CRISPR/Cas9/gRNA solution for the targeted removal of rRNA sequence was prepared and applied to ribo-seq libraries following Mito et al.^35^.

### Data processing

#### Sequencing read processing and mapping

Sequence reads were processed and mapped to the human genome following a modified procedure used by Ingolia et al.^33^. Before aligning to the genome, the adapter and polyA sequences as well as 4 nucleotides at the 5’ end were removed from each read using FASTX-Toolkit. Processed reads mapped to a reference FASTA file composed of human rRNA, tRNA and snoRNA sequences were removed. The remaining sequence reads were mapped to the GRCh38 human genome using TopHat2^36^ with splice-junction information from GENCODE GTF (release 37). The mapping procedure allowed a maximum of 2 mismatches and only uniquely mapped reads were retained. Levels of protein translation were estimated by counting the number of ribosome profiling reads aligning to each gene based on GENCODE annotation using featureCounts^37^.

#### Data filtering and transformation

To focus our analysis on sufficiently quantitated genes, we only analyzed genes containing at least one sequencing read from each sample, which resulted in a dataset including 12,357 genes (Data S1). For spike-in oligomers, all available quantifications were included in the analyses. Unless otherwise specified, before downstream analyses, count data were log2 transformed after the addition of 0.25 pseudo-count to avoid creating singular values in log scale.

### Analyses

Statistical analyses were performed in the R statistical computing environment (version 4.0.4). Linear regression analyses were performed using the lm() function. Spearman and Pearson correlations and corresponding tests were performed using cor() and cor.test(). Student’s t-Test were performed using t.test(). Wilcoxon Rank Sum tests were performed using wilcox.test().

#### Normalization

TMM normalization was performed using the cpm() function from the edgeR package^38^ with effective library size calculated using the calcNormFactors() function. Default parameters were used for the trimmed mean calculation.

RUVg normalization was performed using the RUVSeq package^1^. Factor analysis for identifying unwanted variables was performed by applying the RUVg() function on raw ribo-seq counts using the default settings with the spike-in oligomers as control genes. Note that the factor analysis was done on count data after filtering out genes that do not have at least one count per sample, but without the commonly applied preprocessing of upper quartile normalization, which could remove biologically meaningful coverage differences resulting from the stress treatment. RUVg normalized counts were extracted using the normCounts() function for downstream analyses to visualize the impact of normalization.

The mitoTMM normalization was performed using the same procedure as a regular TMM normalization except the effective library size is calculated using mitochondrial ribosome footprint counts. Mitochondrial ribosome footprints were defined as ribosome protected fragments that were mapped to GENCODE transcripts (release 37) on chrM.

Respective quantile normalization was performed by applying normalizeQuantiles() function from the limma package^39^ separately to treated and control samples.

#### Differential expression tests

Differential expression analyses were performed using the limma package^39^. After processing with various normalization and transformation procedures, functions lmFit() and eBayes() were applied sequentially to perform differential expression tests under a linear modeling framework. When testing for treatment effect for endogenous genes, both library construction batch and donor identity were fitted as covariate in the linear model. More specifically, for each gene, expression level E across sample j is modeled by the treatment effect T as the predictor and the library batch effect B and the cell line effect C as covariates in the following equation:

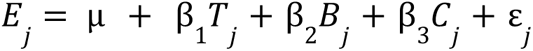

Where μ is the intercept term and ε is the residual. To identify differentially expressed genes, we tested the null hypothesis of β_1_ is zero using the empirical Bayes moderated t-statistics from the eBayes() function. False discovery rates were calculated using the Benjamini-Hochberg procedure.

## Authors’ contributions

SHW, SKB conceived and designed the study. SKB performed the experiments. AWS,SHW performed the majority of data analysis. ZZ, SKB, SHW designed the panel of spike-in oligomer controls. SHW wrote the paper. SHW, AWS, SKB, ZZ commented on and revised the paper.

## Acknowledgements

We thank members of the Wang lab for helpful discussions. The University of Chicago Genomics Facility for collecting the sequencing data.

