## Supplemental Figures for "Transcriptome-wide profiling of acute stress induced changes in ribosome occupancy level using external standards"

Supplemental Tables & Figures

Table S1 - Spike-in Oligomers

| name | RNA_Sequence |
|---|---|
| **Batch 1** | |
| A1 | AACGCCCAGAAACGCCCCGACAAGGCGG |
| A2 | AAAUAACCACGAAGAGCCCAAGAGAUUA |
| A3 | GUUCGACACAGGGGAAAGGGGGGAGAGG |
| A4 | CAUGACACACAAGGGGACAAAAGGUCGG |
| B1 | CAUACAAAAAGAGGGAAGCAAGGGUAGA |
| B2 | UACCGCAAGACAGGAACCGAGAGAGUUA |
| B3 | GUUUCCCCAGAAGAGACCCGCCGCGACC |
| B4 | GUCAGACAAGAGGGCAGAAGAGAGAUAG |
| **Batch 2** | |
| C1 | CAGGGAAAACACGAAGGAGCAGAGGUUU |
| C2 | UAAAGCAAACAAGAACAAGGCCGAUGGG |
| C3 | UUGACGAAACCGAGACCCCCCCAGUAAC |
| C4 | UGUUAGACGCAAGGAAAACACGAGACAC |
| D1 | CAAGACAAGAACAGCCGGGACAAAAUAG |
| D2 | AAACAGACGACCGCAGGACCCAAGCGUG |
| D3 | ACAAAGAGACCCCGCAGAGCCCACAGAG |
| D4 | CCUUACAGAGAGGGGAAGGGAACGUUCC |

Table S2 - Mapping Statistics

| sample | seq_depth | unique_reads | spikein_reads | rRNA_reads |
|---|---|---|---|---|
| C193T | 50382005 | 2016717 | 31190 | 40308561 |
| C193U | 85058780 | 3193118 | 49737 | 69150869 |
| C204T | 57025679 | 3010568 | 18164 | 46414163 |
| C204U | 78143644 | 3544299 | 21620 | 63399018 |
| C505T | 48459128 | 2199813 | 7519 | 39707227 |
| C505U | 98746922 | 4167985 | 14694 | 80827933 |
| E193T | 51904657 | 744639 | 4935 | 44043792 |
| E193U | 136927240 | 1799231 | 12670 | 116181233 |
| E204T | 90395514 | 1248240 | 98469 | 77373538 |
| E204U | 87479545 | 1141768 | 90511 | 74942324 |
| E505T | 41650935 | 649571 | 17792 | 35208261 |
| E505U | 47206608 | 702012 | 18485 | 40547952 |
| **Total** | **873380657** | **24417961** | **385786** | **728104871** |
| **average** | **72781721** | **2034830** | **32149** | **60675406** |

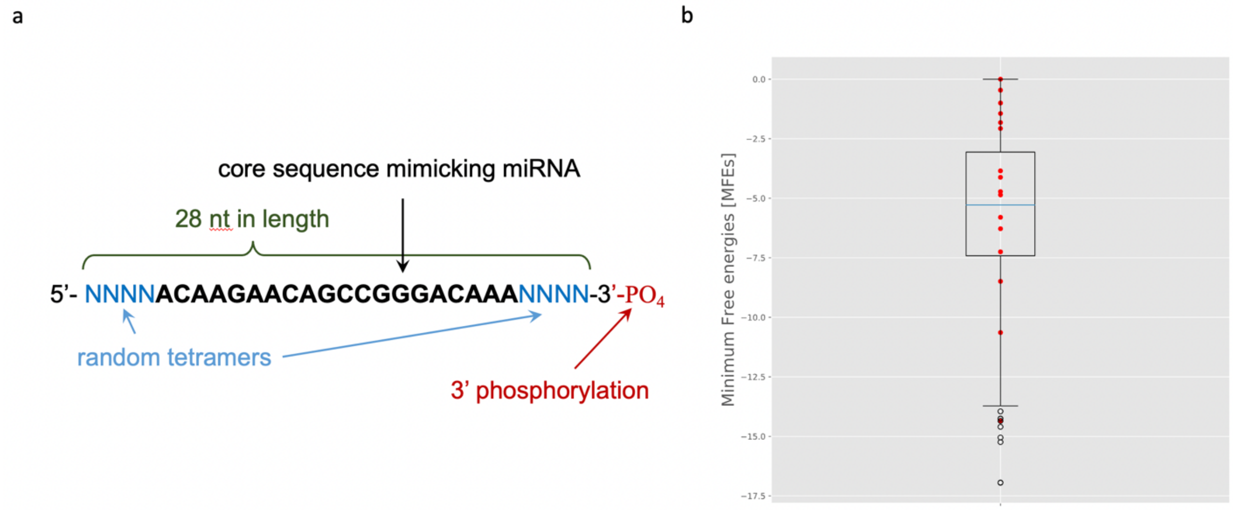


Figure S1. Spike-in oligomer design. (a) Diagram illustrating key features of spike-in oligomers. (b) Minimum free energy profile of 16 spike-in oligomers used in this study (red data points) plotted on top of a boxplot summarizing the minimum free energy profile of endogenous human miRNAs. The maximum and minimum values for the boxplot (i.e. the whiskers) are defined by the miRNAs with minimum free energy values closest to (but without exceeding) 1.5 times of the interquartile range extending from the box.

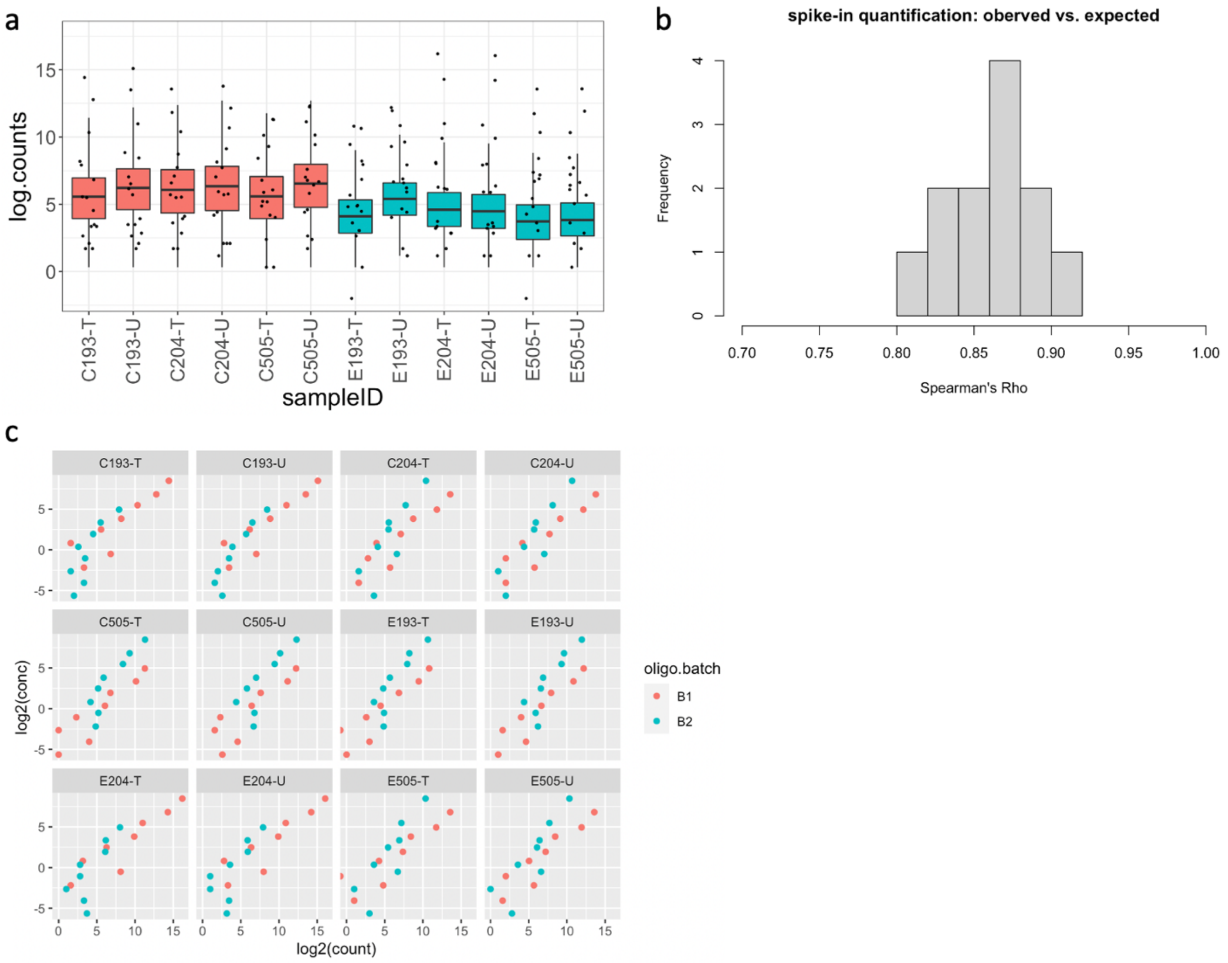


Figure S2. Quantitative ranges and the correlations between the observed and the expected quantifications for the 16 spike-in oligomers. (a) Boxplots summarizing the overall distribution of ribosome occupancy level for all analyzed genes overlaid with data points indicating the quantification level of each of the 16 spike-in oligomers. Boxes are color coded to distinguish control samples (red) from Sodium Arsenite treated samples (blue). The maximum and minimum values for the boxplot (i.e. the whiskers) are defined by the genes with quantification levels closest to (but without exceeding) 1.5 times of the interquartile range extending from the box. (b) Histogram summarizing the distribution of Spearman's Rho between the observed (log2 counts) and the expected (log2 concentration) across 12 samples. (c) Quantifications of spike-in oligomers (log2 counts) plotted against the expected (log2 concentration) for each individual sample. Each data point represents a spike-in oligomer and the color code indicates the manufacturing batch. Individual panels are labeled with abbreviated sample information (e.g. C505-T indicates a control sample [C] of cell line GM18505 [505] from the library preparation batch T [T]).

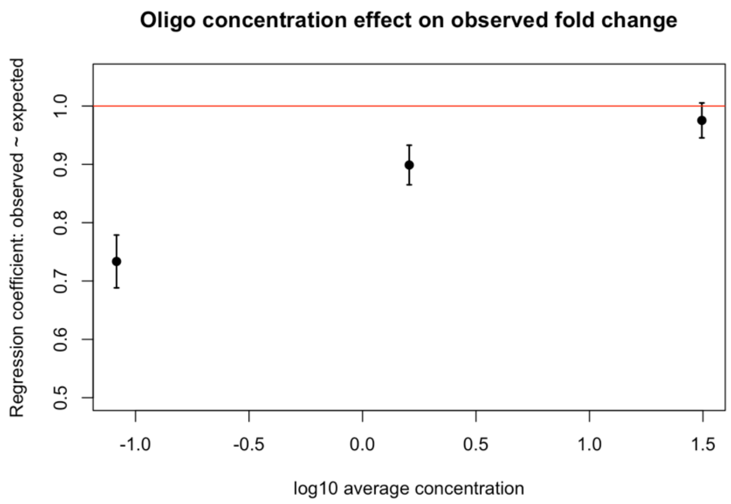


Figure S3. Quantifications for spike-in oligomers used in lower concentrations show lower correlations to the expected. Oligomers were stratified into 3 concentration strata and the average concentration (x-axis) for each stratum is plotted against the regression coefficient calculated from using the expected fold change as predictor for the observed. Red horizontal line marks the slope of 1. Error bars represent the standard error of regression coefficients from each concentration stratum.

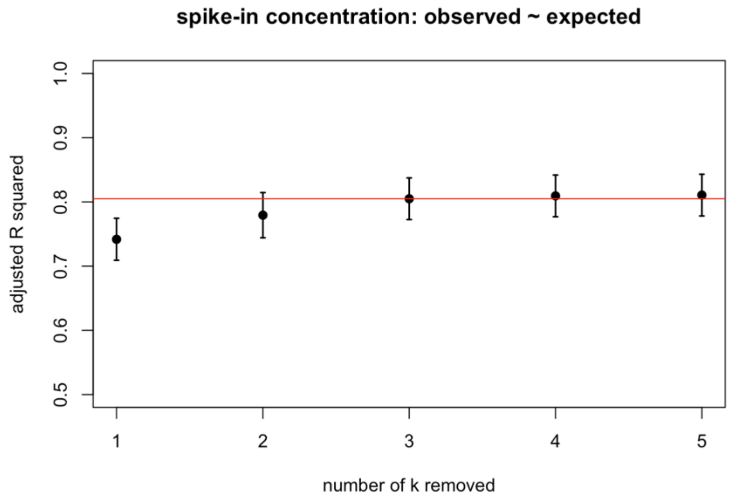


Figure S4. Evaluating the impact of spike-in based RUVg normalization on subsampled spike-in quantifications. Spike-in oligomers were randomly divided into two equal size groups and one group was used as control genes for estimating hidden unwanted variables (k) while the other group was used to visualize the impact of incrementally removing unwanted variables on the correlation. R-squared of a linear regression fit between the observed and expected spike-in quantifications were plotted against the number of k removed. The error bars represent the standard error of R-squared across 10 iterations of random subsampling. The red horizontal line marks the mean R-squared resulted from removing 3 unwanted variables (i.e. k = 3).

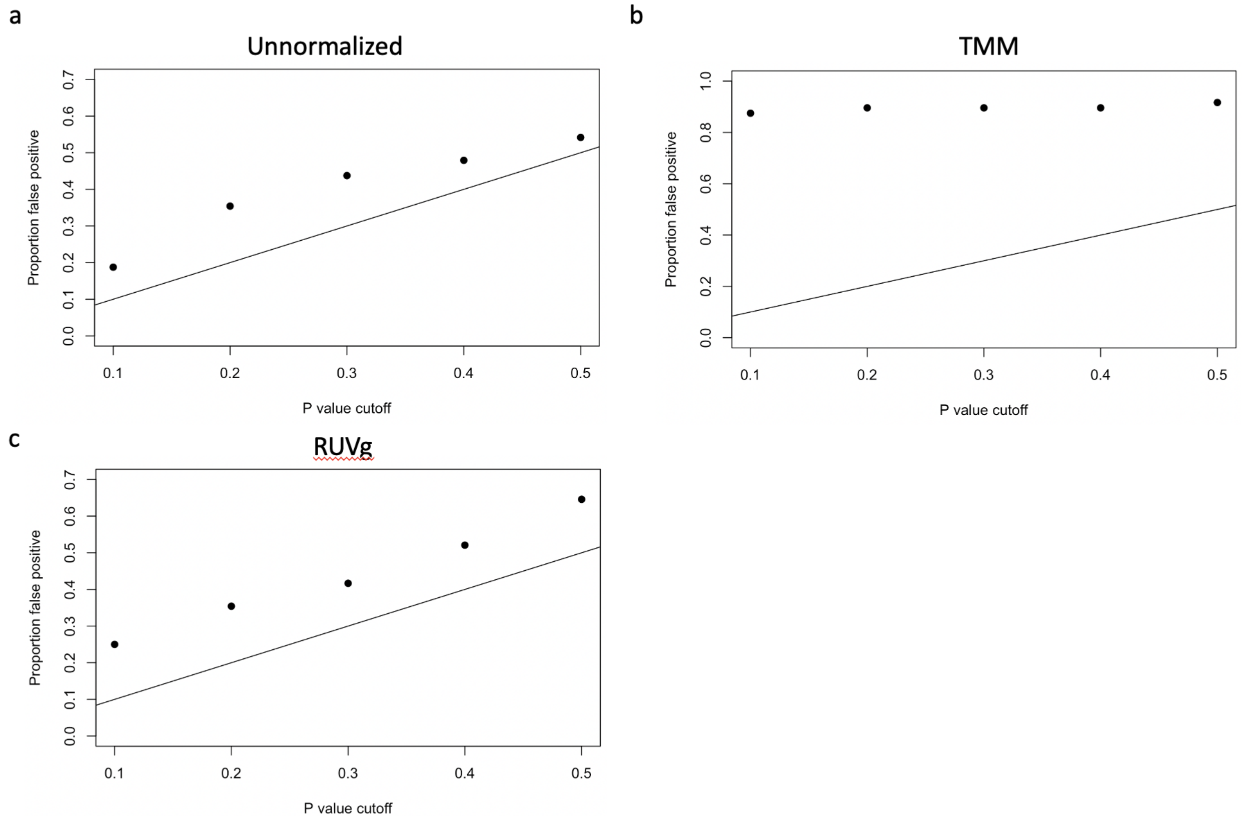


Figure S5. Proportion of significant differences found in spike-in constructed null comparisons across a range of P value cutoffs. The black line marks the expected proportion of false positives. (a) log2 transformed count data (b) TMM normalized log2 count (c) RUVg normalized log2 count (using spike-in oligomers as control genes).

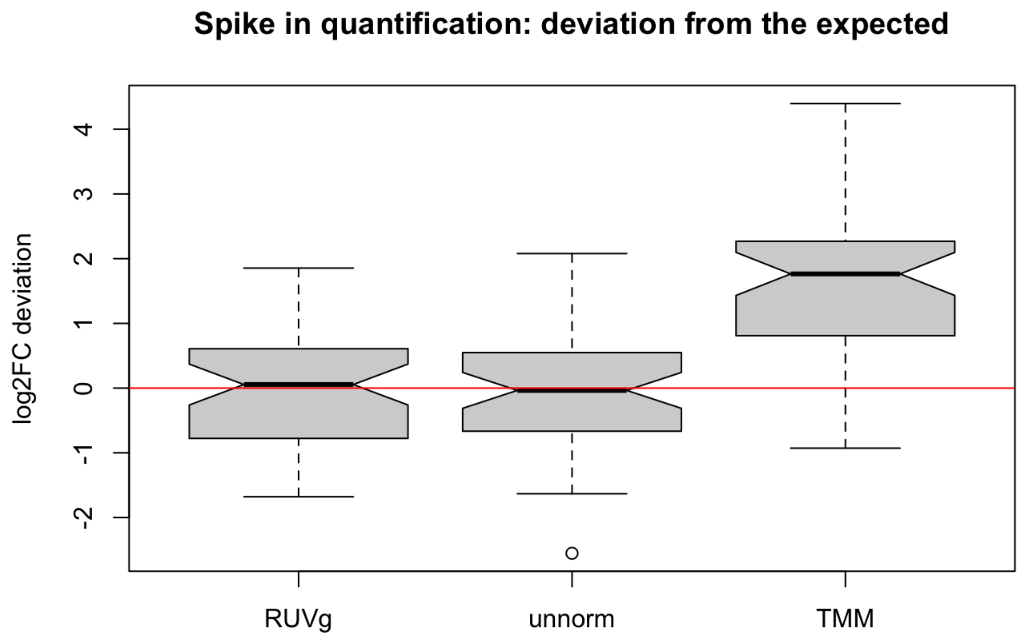


Figure S6. Boxplots comparing deviations from the expected fold change for spike-in constructed true positives between results from spike-in based RUVg normalization (RUVg), without normalization (unnorm) and TMM normalization (TMM). The maximum and minimum values (i.e. the whiskers) are defined by the data point with value closest to (but without exceeding) 1.5 times of the interquartile range extending from the box. The red horizontal line marks the expected 0 deviation.

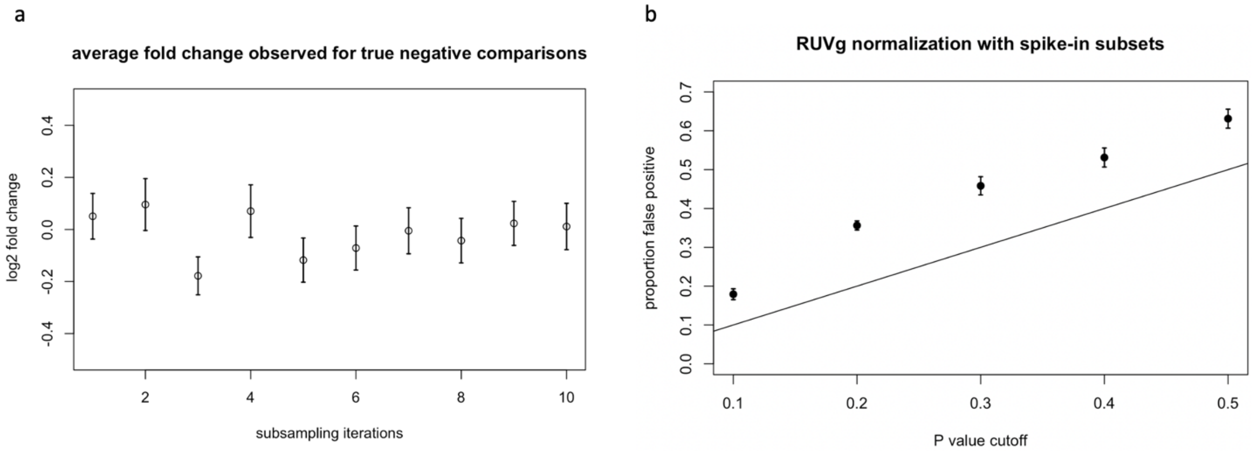


Figure S7. Observed fold change and proportion false positives from spike-in constructed null comparisons across 10 iterations of random subsampling. (a) Observed fold change, error bars represent the standard error across null comparisons, which were constructed using test group oligomers. (b) Proportion false positives, error bars represent the standard error across 10 iterations of random subsampling. The black line marks the expected proportion of false positives.

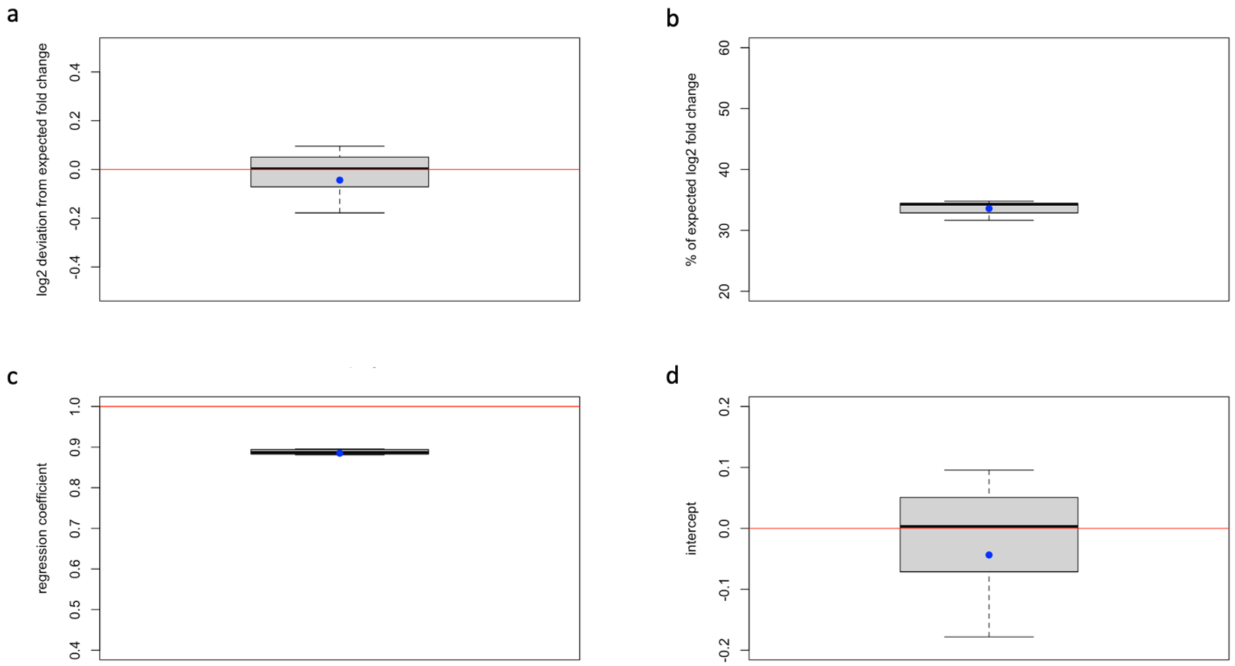


Figure S8. Boxplots summarizing the range of values observed from spike-in constructed true positives across 10 iterations of random subsampling. (a) Deviation from the expected fold change (b) Deviation from the expected fold change standardized by effect size (c) Regression coefficient from using the expected fold change as the predictor (d) Intercept calculated from using the expected fold change as the predictor. The red horizontal lines mark the ideal value. The blue data point marks the value calculated from using the full set of spike-in oligomers.

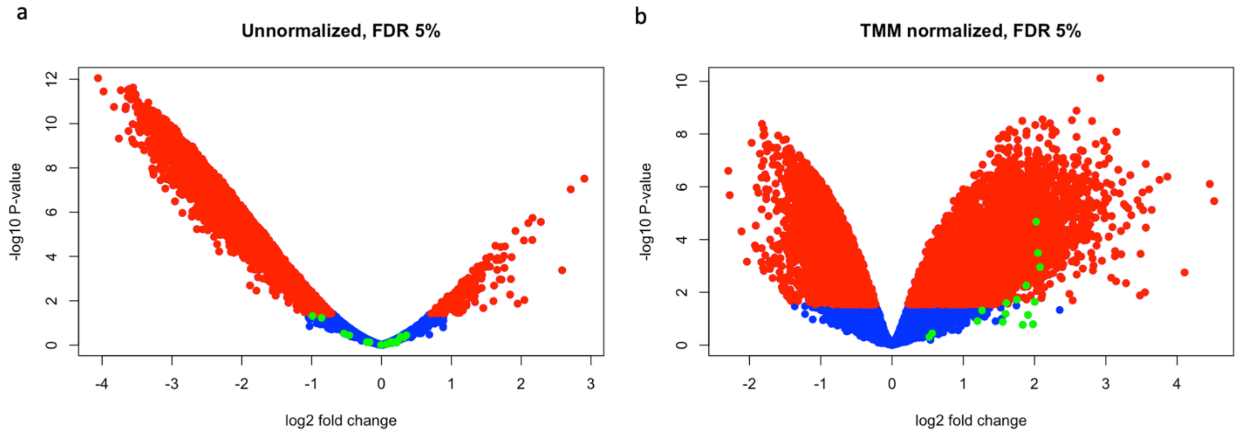


Figure S9. Volcano plots showing the relationship between fold change (treatment versus control) and p value from differential expression tests for (a) Data without normalization (unnormalized) and (b) TMM normalized data. Data points are color-coded in green for spike-in oligomers, in red for endogenous genes that are significantly differentially expressed at 5% FDR, and in blue for endogenous genes that are not differentially expressed.

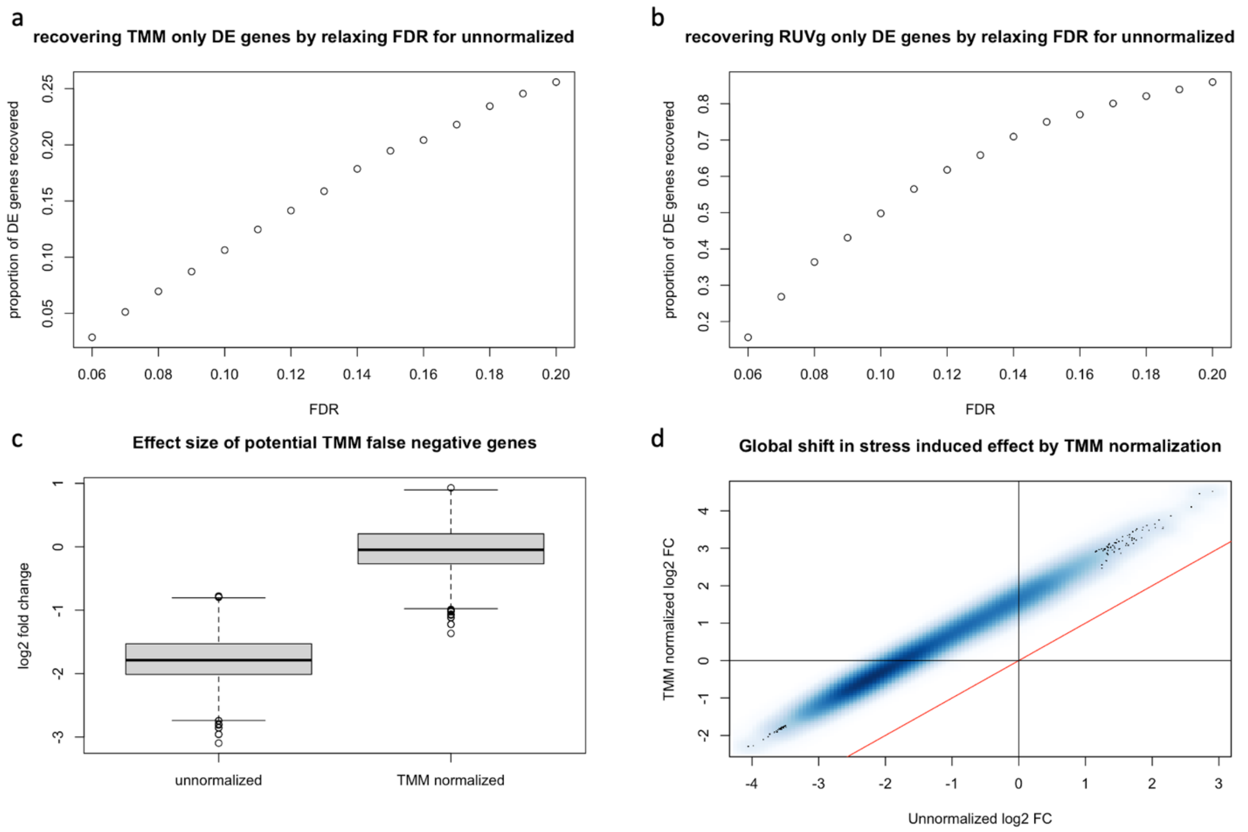


Figure S10. Impact of TMM normalization on stress response ribo-seq data. (a,b) Proportion of differentially expressed genes found only in normalized data that were recovered by relaxing significance cutoffs (x-axis) for differential expression tests performed using data without normalization. Note the low recovery rate from TMM normalized data (a) in contrast to the high recovery rate from spike-in based RUVg normalized data (b). (c) Boxplots summarizing effect size change from TMM normalization for differentially expressed genes found in unnormalized data but not in TMM normalized data. The maximum and minimum values (i.e. the whiskers) are defined by the data point with value closest to (but without exceeding) 1.5 times of the interquartile range extending from the box. (d) A smoothed color density representation of a scatter plot showing the impact of TMM normalization on stress induced translational changes. All endogenous genes deemed sufficiently quantitated are included in the plot with color density proportional to the number of genes presented in each unit region (bin) of the plot. Black lines mark 0 fold change genes for each dataset. Red line marks the perfect correlation (i.e. an intercept of 0 and a slope of 1)

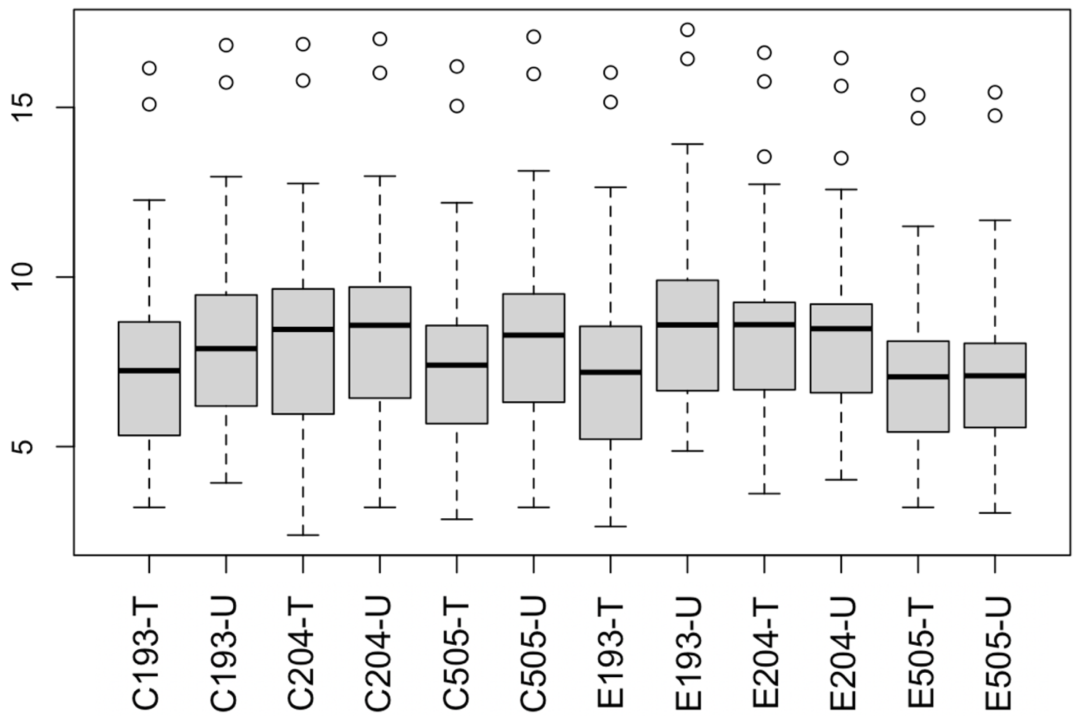


Figure S11. Boxplots summarizing the distribution of mitochondrial ribosome footprint counts across samples. The maximum and minimum values (i.e. the whiskers) are defined by the data point with value closest to (but without exceeding) 1.5 times of the interquartile range extending from the box.

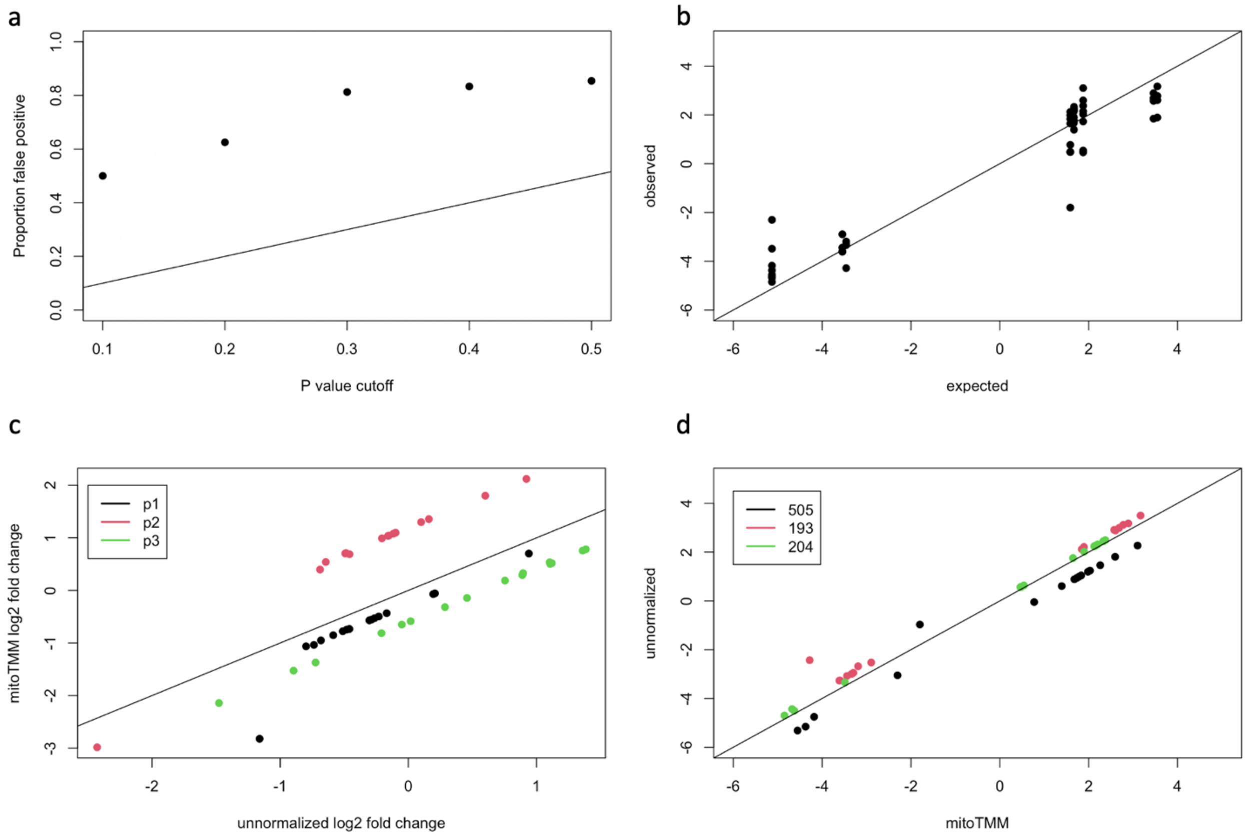


Figure S12. Impact of mitoTMM normalization on spike-in oligomer quantification. (a) Proportion of false positives found from spike-in constructed nulls in mitoTMM normalized data across a wide range of significance levels. (b) Observed log2 fold change for spike-in constructed true positives in mitoTMM normalized data plotted against the expected. (c) Observed fold change for spike-in constructed nulls (between individual by pool comparisons) in mitoTMM normalized versus unnormalized data. Data points are color-coded by spike-in pool designation. (d) Observed fold change for spike-in constructed true positives (between pool by individual comparisons) in mitoTMM normalized versus unnormalized data. Data points are color-coded by individual identification (i.e. the origin of cell lines).

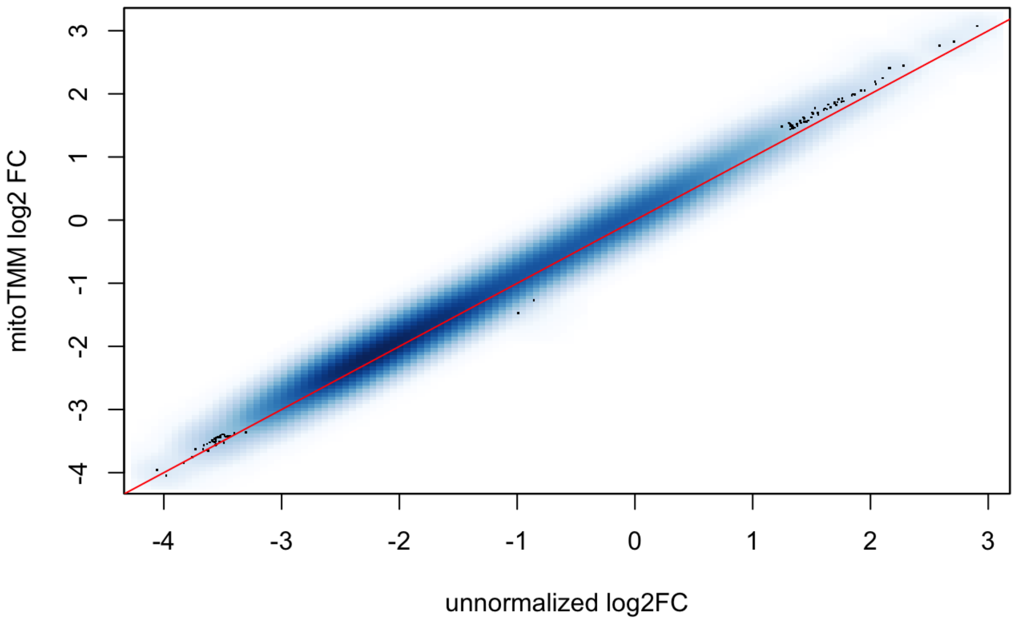


Figure S13. Impact of mitoTMM normalization on quantification of endogenous genes. Log2 fold change between treatment and control samples are visualized in a smoothed color density representation of a scatter plot. All endogenous genes deemed sufficiently quantitated are included in the plot with color density proportional to the number of genes presented in each unit region (bin) of the plot. Red line marks the perfect correlation (i.e. an intercept of 0 and a slope of 1).

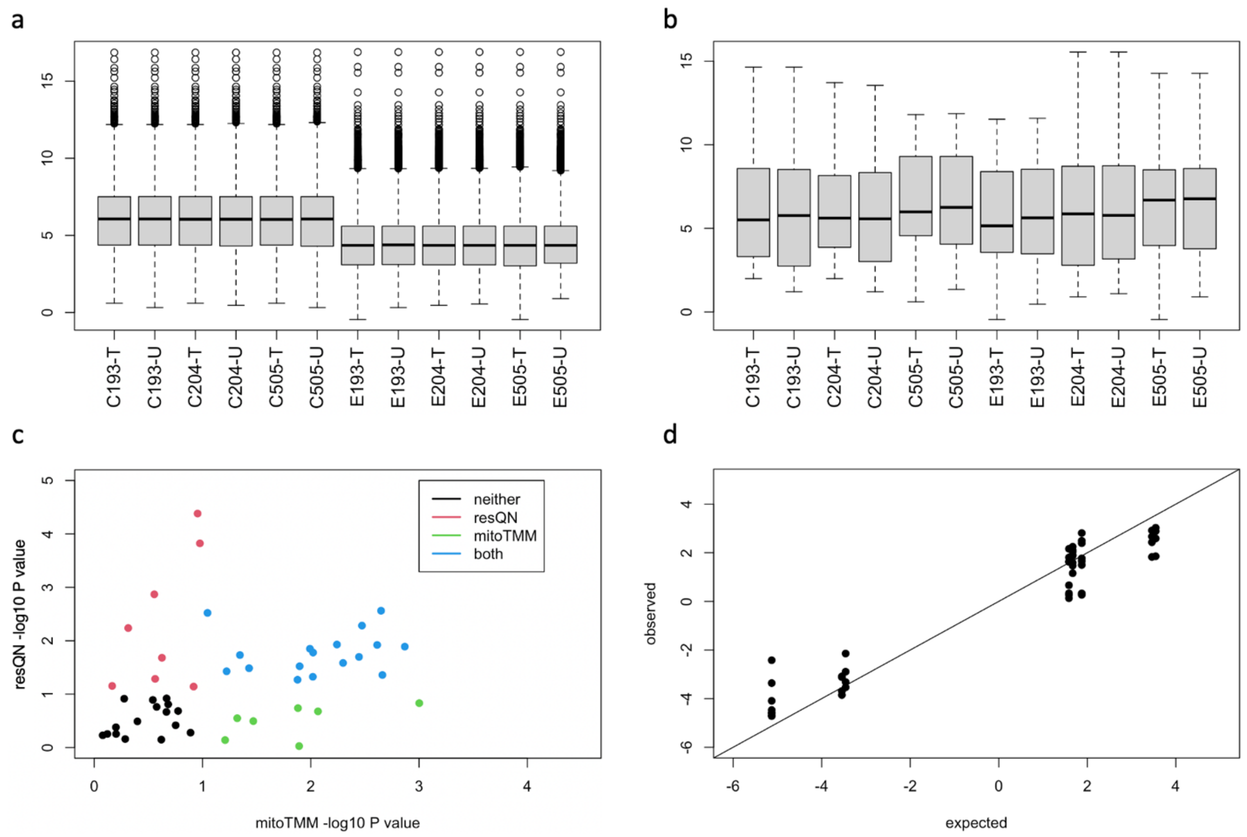


Figure S14. Impact of respective quantile normalization on expression quantification. (a,b) Boxplots summarizing overall distributions of expression quantifications for endogenous genes (a) and for spike-in oligomers (b) across samples after respective quantile normalization. The maximum and minimum values (i.e. the whiskers) are defined by the data point with value closest to (but without exceeding) 1.5 times of the interquartile range extending from the box. (c) Comparing false positives found for spike-in constructed nulls in respectively quantile normalized data versus mitoTMM normalized data. Data points are color-coded by significance ($P < 0.1$). Key for the color code presented in the inset; resQN stands for respective quantile normalization. (d) Observed log2 fold change for spike-in constructed true positives from respectively quantile normalized data plotted against the expected.

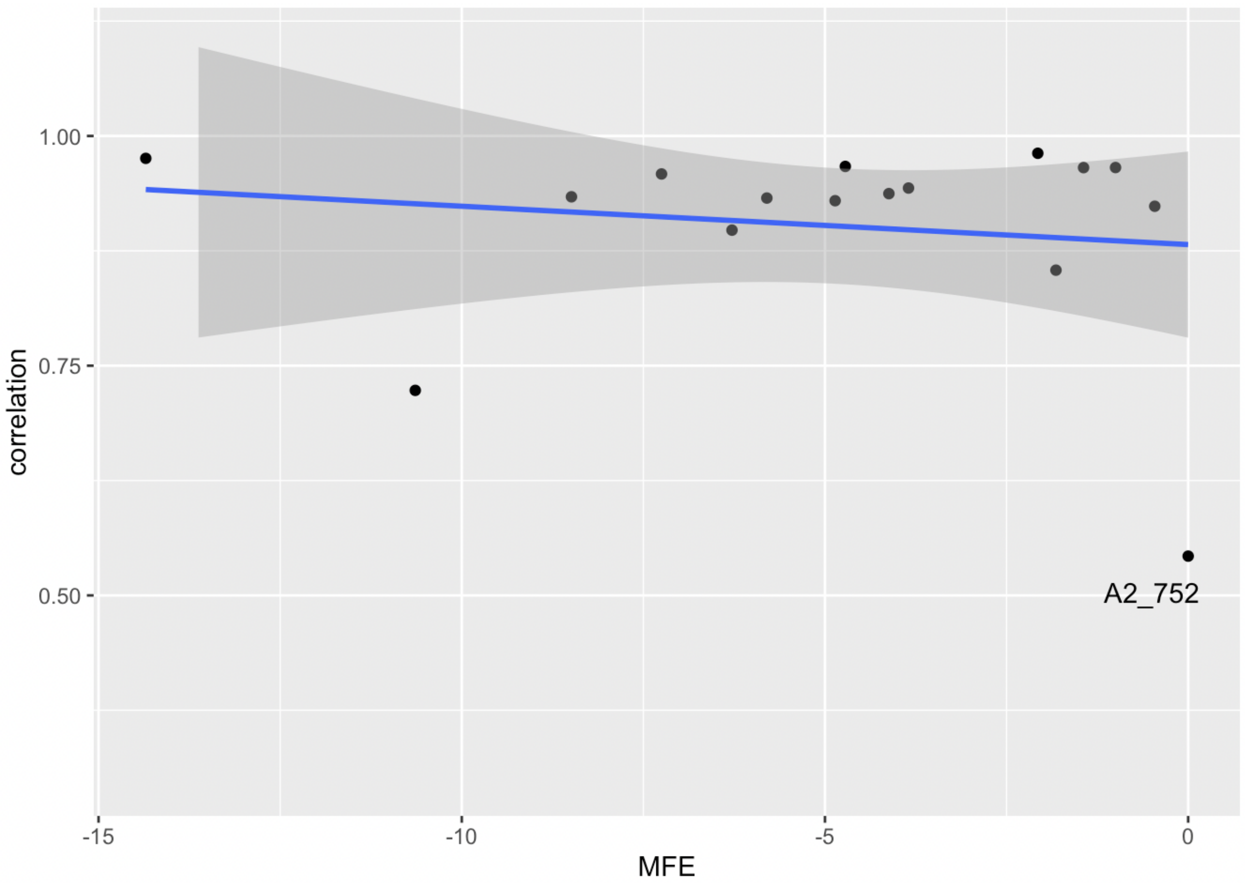


Figure S15. A scatter plot visualizing the association between spike-in oligomer Minimum Free Energy (MFE) and the correlations between the observed and the expected quantification for spike-in oligomers. The blue trend line and the corresponding grey shaded area represents the regression coefficient of a linear model fit using MFE as the predictor and the corresponding 95% confidence intervals.

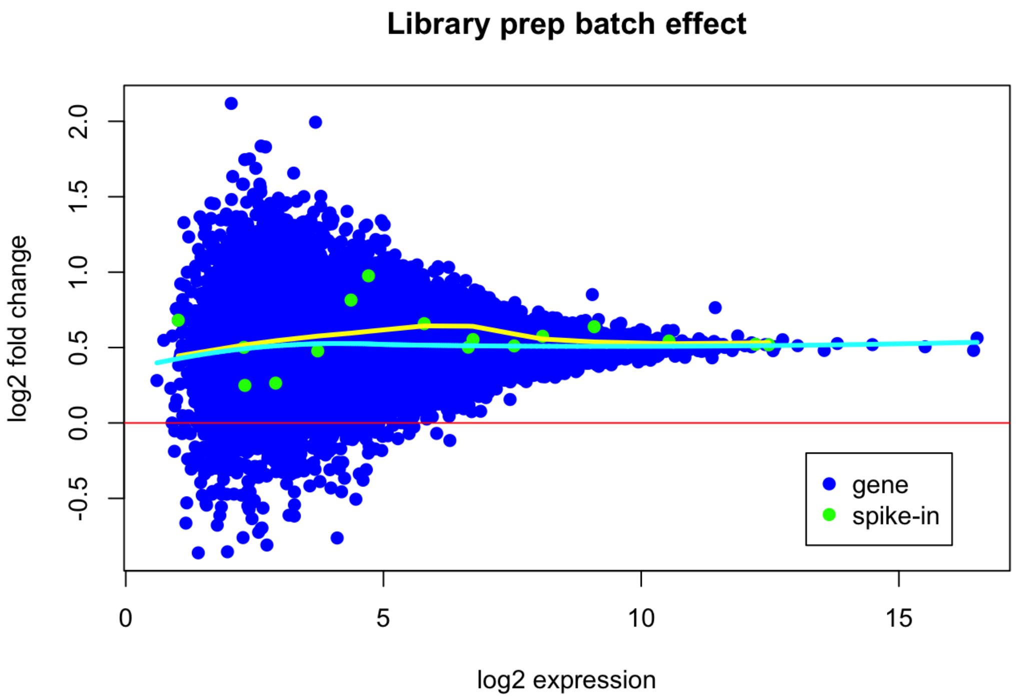


Figure S16. An M-A plot visualizing impact of library preparation batch effect on the quantification of endogenous genes and spike-in oligomers. Color code for data points are presented in the inset. Red horizontal line marks no differences between library preparation batches; loess trend lines were calculated separately for endogenous genes (cyan) and for spike-in oligomers (yellow) using an α of 95%.
